# Extended field-of-view ultrathin microendoscopes for high-resolution two-photon imaging with minimal invasiveness in awake mice

**DOI:** 10.1101/2020.05.14.095299

**Authors:** Andrea Antonini, Andrea Sattin, Monica Moroni, Serena Bovetti, Claudio Moretti, Francesca Succol, Angelo Forli, Dania Vecchia, Vijayakumar P. Rajamanickam, Andrea Bertoncini, Stefano Panzeri, Carlo Liberale, Tommaso Fellin

**Affiliations:** Optical Approaches to Brain Function Laboratory, Istituto Italiano di Tecnologia, 16163 Genova, Italy; Nanostructures Department, Istituto Italiano di Tecnologia, 16163 Genova, Italy; University of Genova, 16126 Genova, Italy; Neural Coding Laboratory, Istituto Italiano di Tecnologia, 16163 Genova and 38068 Rovereto, Italy; Neural Computation Laboratory, Center for Neuroscience and Cognitive Systems @UniTn, Istituto Italiano di Tecnologia, 38068 Rovereto, Italy; Center for Mind and Brain Sciences (CIMeC), University of Trento, 38068 Rovereto, Italy; Biological and Environmental Sciences and Engineering Division (BESE), King Abdullah University of Science and Technology (KAUST), Thuwal 23955-6900, Saudi Arabia

## Abstract

Imaging neuronal activity with high and homogeneous spatial resolution across the field-of-view (FOV) and limited invasiveness in deep brain regions is fundamental for the progress of neuroscience, yet is a major technical challenge. We achieved this goal by correcting optical aberrations in gradient index lens-based ultrathin (≤ 500 µm) microendoscopes using aspheric microlenses generated through 3D-microprinting. Corrected microendoscopes had extended FOV (*eFOV*) with homogeneous spatial resolution for two-photon fluorescence imaging and required no modification of the optical set-up. Synthetic calcium imaging data showed that, compared to uncorrected endoscopes, *eFOV*-microendoscopes led to improved signal-to-noise ratio and more precise evaluation of correlated neuronal activity. We experimentally validated these predictions in awake head-fixed mice. Moreover, using *eFOV-*microendoscopes we demonstrated cell-specific encoding of behavioral state-dependent information in distributed functional subnetworks in a primary somatosensory thalamic nucleus. *eFOV-*microendoscopes are, therefore, small-cross-section ready-to-use tools for deep two-photon functional imaging with unprecedentedly high and homogeneous spatial resolution.

## Introduction

The amount of information carried by neural ensembles and the impact that ensemble activity has on signal propagation across the nervous system and on behavior critically depend on both the information and tuning properties of each individual neurons and on the structure of correlated activity, either at the level of correlations between each pair of neurons or at the whole network level ^1-5^. To study neuronal population coding, it is thus essential to be able to measure with high precision, large signal-to-noise-ratio (SNR), and without distortions the activity of individual neurons and the relationship between them ^6^. Two-photon imaging makes it possible to record the activity of many hundreds of individual neurons simultaneously and provides reliable measures of correlated neuronal events ^7-11^. Light scattering within the brain, however, strongly affects the propagation of excitation and emission photons, making effective imaging increasingly difficult with tissue depth ^12-14^. Various strategies have been developed to improve imaging depth in multi-photon fluorescence microscopy ^15-22^, allowing the visualization of regions 1-1.6 mm below the brain surface. However, deeper imaging requires the use of implantable microendoscopic probes or chronic windows, which allow optical investigation of neural circuits in brain regions that would otherwise remain inaccessible ^23-27^.

A critical barrier to progress is the lack of availability of microendoscopic devices with small cross-sections that maintain cellular resolution across a large FOV, to allow high-resolution and high SNR two-photon population imaging on a large number of neurons while minimizing tissue damage. Current microendoscopes for deep imaging are frequently based on grade index (GRIN) rod lenses, which typically have diameter between 0.35-1.5 mm and are characterized by intrinsic optical aberrations ^25^. These aberrations are detrimental in two-photon imaging because they decrease the spatial resolution and lower the excitation efficiency, especially in the peripheral parts of the FOV. This leads to uneven SNR and spatial resolution across the FOV and therefore restricted effective FOV ^28, 29^. This is a serious issue when measuring correlated neuronal activity, because signals sampled at the periphery of the FOV will be more contaminated by neuropil or by activity of neighboring cells compared to signals sampled near the optical axis. Optical aberrations in GRIN microendoscopes can be corrected with adaptive optics which, however, requires significant modification of the optical path ^28, 30, 31^ and may limit the temporal resolution of functional imaging over large FOVs ^28^. Alternatively, the combination of GRIN lenses of specific design with plano-convex lenses within the same microendoscopic probe has been used to increase the Numerical Aperture (NA) and to correct for aberrations on the optical axis ^25^. However, technical limitations in manufacturing high-precision free-form optics with small lateral dimensions have so far prevented improvements in the performances of GRIN microendoscopes with lateral diameter < 1 mm using corrective optical microelements ^32^.

Here, we report the design, development, and characterization of a new approach to correct aberrations and extend the FOV in ultrathin GRIN-based endoscopes using aspheric lenses microfabricated with 3D micro-printing based on two-photon lithography (TPL) ^33^. These new endoscopic probes are ready-to-use and they require no modification of the optical set-up. We applied *eFOV*-microendoscopes to study the encoding of behavioral state-dependent information in a primary somatosensory thalamic nucleus of awake mice in combination with genetically encoded calcium indicators.

## Results

### Optical simulation of eFOV-microendoscopes

Four types (type I-IV) of *eFOV*-microendoscopes of various length and cross section were designed, all composed of a GRIN rod, a glass coverslip and a microfabricated corrective aspheric lens (Fig. 1). One end of the GRIN rod was directly in contact with one side of the glass coverslip while the corrective aspheric lens was placed on the other side of the coverslip. GRIN rods and corrective lenses were different in each of the four types of *eFOV*-microendoscopes (lateral diameter, 0.35-0.5 mm; length, 1.1-4.1 mm; all 0.5 NA). The glass coverslip was 100 µm thick for type I, III, IV *eFOV*-microendoscopes and 200 µm thick for type II *eFOV*-microendoscopes. This design did not require additional cannulas or clamping devices ^34, 35^ that would increase the lateral size of the microendoscope assembly or reduce its usable length, respectively.

**Fig. 1.**
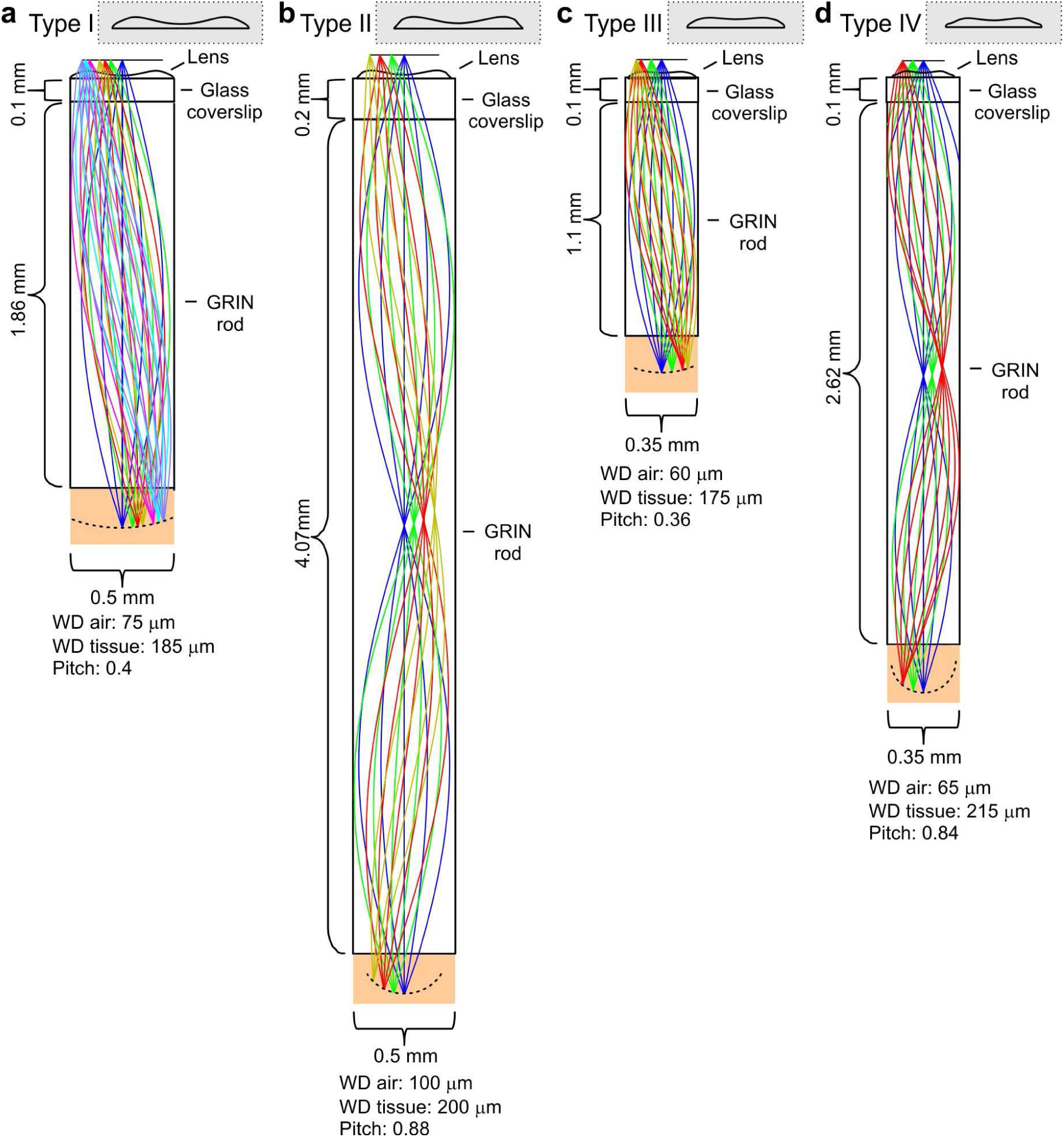
Optical design of *eFOV*-microendoscopes. **a-d**, Ray trace simulations for the four different *eFOV*-microendoscopes (type I-IV). The insets show the profiles of corrective polymeric lenses used in the different *eFOV*-microendoscopes. For each *eFOV*-microendoscope, it is specified the thickness of the coverslip, the length, the diameter of the GRIN rod, the pitch of the GRIN rod, and the working distance, in air or in tissue, at which the simulated correction of aberrations was performed. See also Supplementary Table 1.

For each type of GRIN rod used in the *eFOV*-microendoscopes, ray trace simulations determined the lens profile (Fig. 1) that corrected optical aberrations and maximized the FOV (Fig. 2). In the representative case of type I *eFOV*-microendoscopes, the simulated corrective lens had a diameter of 0.5 mm, height < 40 µm and the coefficients used in equation *(1)* (see Methods) to define the lens profile are reported in Supplementary Table 1. For this type of *eFOV*-microendoscope, the simulated point-spread-function (PSF) at incremental radial distances from the optical axis (up to 200 μm) showed that the Strehl ratio of the system was > 80% (i.e., a diffraction-limited condition is achieved according to the Maréchal criterion ^36^) at a distance up to ∼ 165 µm from the optical axis with the corrective lens, while only up to ∼ 70 µm for the same optical system without the corrective lens (Fig. 2a). This improvement led to a ∼ 5 times increase in the area of the diffraction-limited FOV. Fig. 2b-d reports the Strehl ratio for simulated corrected and uncorrected type II-IV *eFOV*-microendoscopes. Simulations showed that the area of the FOV was ∼ 2-9 times larger for these other types of *eFOV*-microendoscopes, compared to microendoscopes without the corrective lens.

**Fig. 2.**
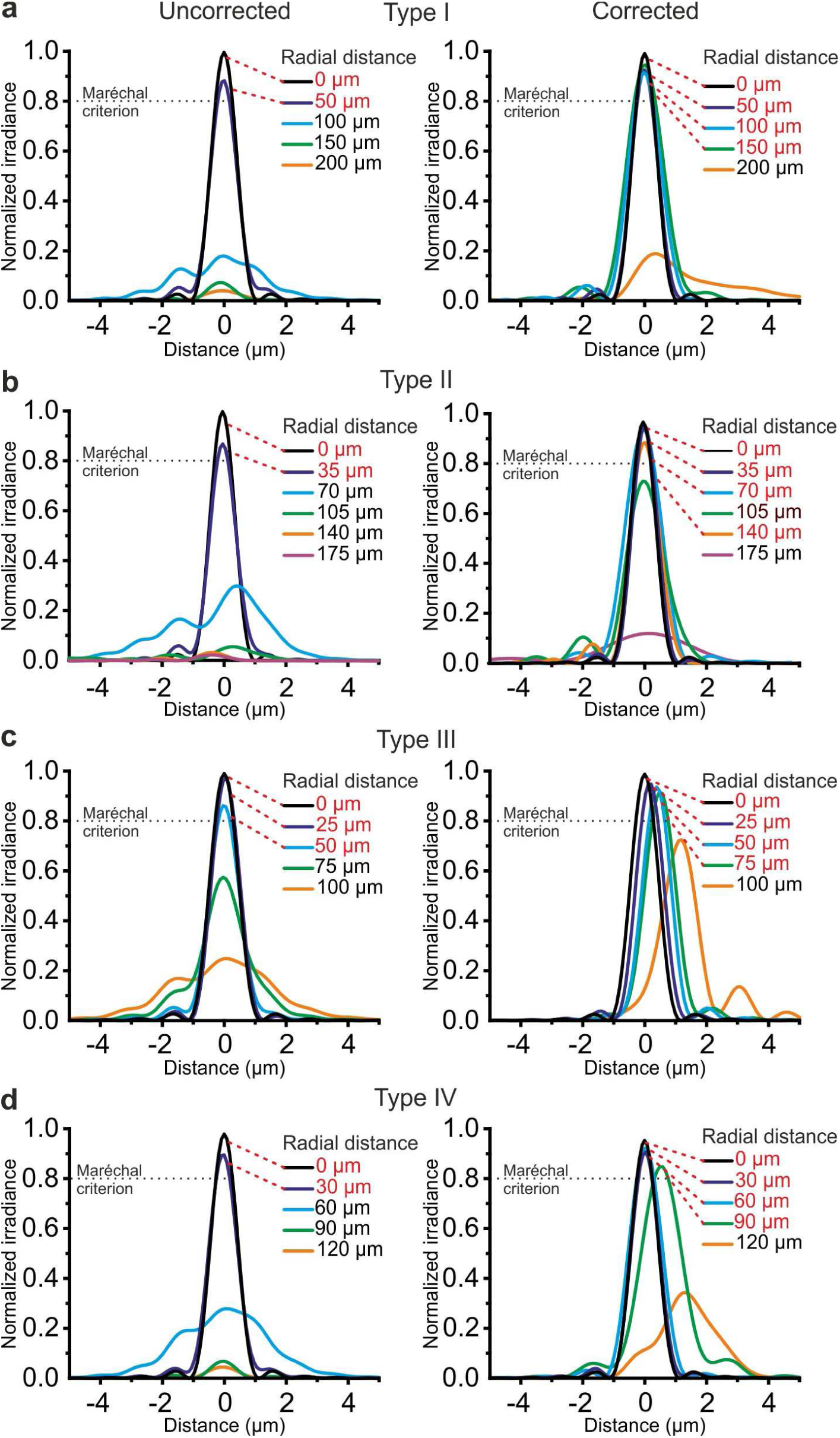
Corrective lenses improve the simulated optical performance of ultrathin microendoscopes. **a**, Simulated diffraction PSFs to assess the Strehl ratio of the designed microendoscope (type I microendoscopes) without the corrective lens (uncorrected, left) and with the corrective lens (corrected, right). PSFs are shown color-coded according to their radial distance from the optical axis. The black dotted line represents the diffraction-limited condition, which was set at 80 % (Maréchal criterion). Distances written in red indicate the radial positions at which the maximal normalized irradiance of the corresponding PSF was > 80 %. **b-d**, Same as in **a** for type II **b**, type III **c**, and type IV **d** microendoscopes.

### Fabrication of eFOV-microendoscopes

Corrective lenses were experimentally fabricated using TPL ^33^(Supplementary Fig. 1a,b) and plastic molding replication ^37^ directly onto the glass coverslip (see Methods). Fabricated lenses had profile largely overlapping with the simulated one (Fig. 1c). The corrective lens was aligned to the appropriate GRIN rod using a customized optomechanical set-up (Supplementary Fig. 2a,b) to generate *eFOV*-microendoscopes. To experimentally validate the optical performance of fabricated *eFOV*-microendoscopes, we first coupled them with a standard two-photon laser scanning system using a customized mount (Fig. 3a,b and Supplementary Fig. 2c,d). We initially measured the on-axis spatial resolution by imaging subresolved fluorescence beads (diameter: 100 nm) at 920 nm. We found that *eFOV*-microendoscopes had similar on-axis axial resolution compared to uncorrected probes (Fig. 3c,e,g,i left and Supplementary Table 2). Given that most aberrations contribute to decrease optical performance off-axis, we repeated the measurements described above and measured the axial and lateral resolution at different radial distances from the center of the FOV. As expected by ray trace simulations (Fig. 2), *eFOV*-microendoscopes displayed higher and more homogeneous axial resolution in a larger portion of the FOV (Fig. 3c,e,g,I left and Supplementary Fig. 3). Defining the effective FOV as that in which FWHM_z_ < 10 µm (dotted line in Fig. 3c,e,g,i left), we found ∼ 3.2 - 9.4 folds larger effective FOV in corrected microendoscopes compared to uncorrected probes (Supplementary Table 2). Importantly, experimental measurements (Fig. 3c,e,g,i) were in good agreement with the prediction of the optical simulations (Fig. 2).

**Fig. 3.**
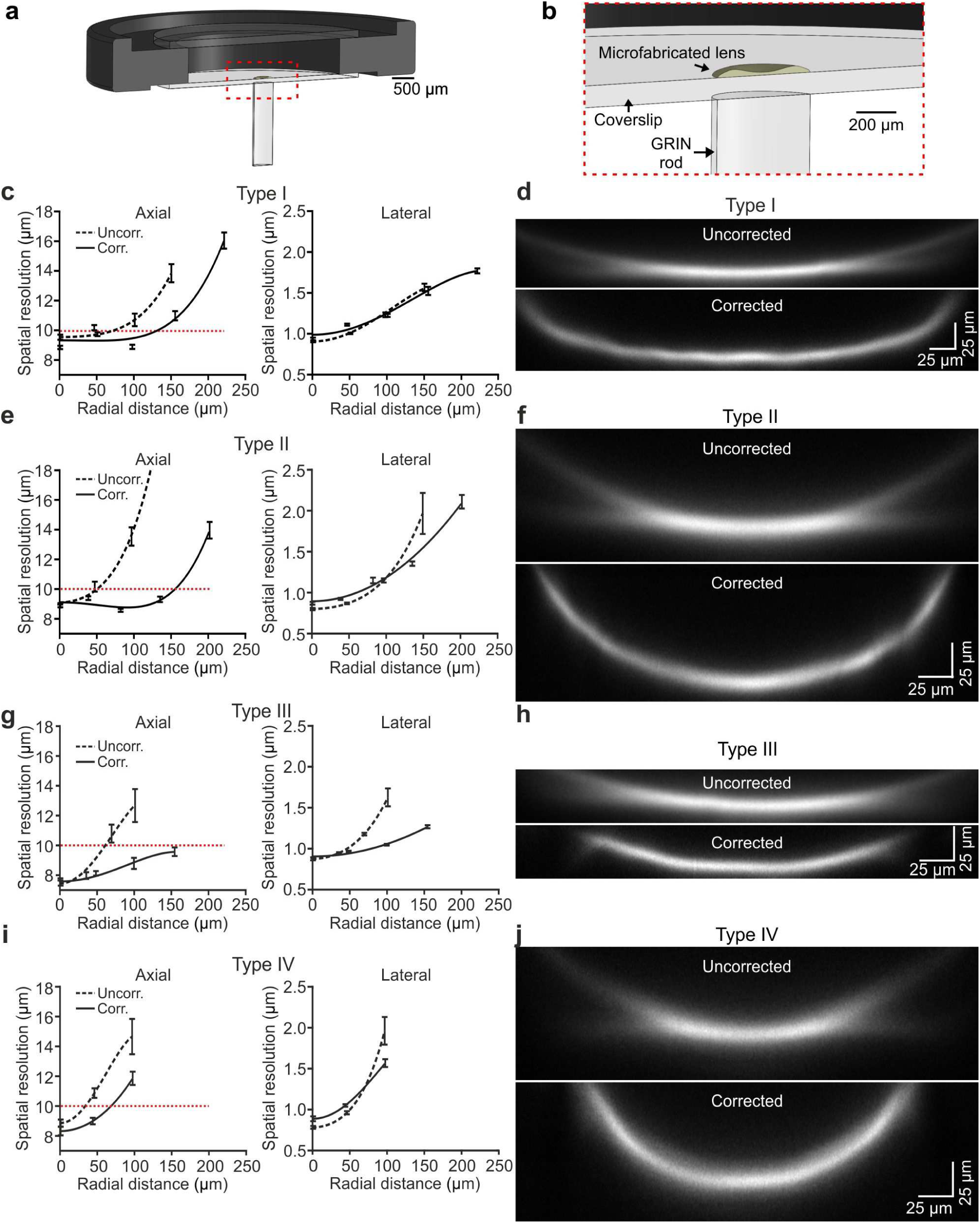
Optical characterization shows enlarged effective FOV in corrected ultrathin microendoscopes. **a**, Schematic of the *eFOV*-microendoscope mount for head implant. The GRIN rod is glued to one side of the glass coverslip, the microfabricated polymer lens to the other side of the coverslip. The coverslip is glued on a circular metal ring that facilitates fixation on the animal’s skull. **b**, Zoom in of the portion highlighted with the red dotted line in **a. c**, Axial (left) and lateral (right) spatial resolution (see STAR Methods for definition) evaluated using subresolved fluorescent beads (diameter: 100 nm) as a function of the radial distance from the center of the FOV for type I uncorrected (dashed line) and corrected (solid line) microendoscopes. Points represent values obtained by averaging at least eight measurements from three different probes (see Supplementary Table 1), while error bar represents standard deviation (sd). Lines are quartic functions fitting the data. The red dashed line indicates a threshold value (10 µm) to define the limit of the effective FOV.**d**, x,z projections (x, horizontal direction; z, vertical direction) of a z-stack of two-photon laser scanning images of a subresolved fluorescent layer (thickness: 300 nm) obtained using a type I *eFOV*-microendoscope, without (uncorrected, top) and with (corrected, bottom) the microfabricated corrective lens. λexc = 920 nm. e,f, Same as in c,d for type II *eFOV*-microendoscopes. **g**,**h**, Same as in **c**,**d** for type III *eFOV*-microendoscopes. **i**,**j**, Same as in c,d for type IV *eFOV*-microendoscopes. See also Supplementary Figs. 1-7 and Supplementary Table 2.

To visualize the profile of fluorescence intensity across the whole diameter of the FOV for both uncorrected and corrected probes, we used a subresolved thin fluorescent layer (thickness: 300 nm) as detailed in ^38^. Fig. 3d,f,h,j shows the x, z projections of the z-stack of the subresolved fluorescent layer for uncorrected and corrected type I-IV microendoscopes. In agreement with the measurements of spatial resolution using subresolved fluorescent beads (Fig. 3c,e,g,i), *eFOV*-microendoscopes displayed higher intensity and smaller FWHM_z_ in peripheral portions of the FOV compared to uncorrected probes (Fig. 3d,f,h,j). *eFOV*-microendoscopes were characterized by a curved FOV and this distortion was evaluated using a fluorescent ruler (Supplementary Fig. 4) and corrected for in all measurements of spatial resolution (Fig. 3 and Supplementary Table 2). The ability of *eFOV*-microendoscopes to image effectively larger FOV compared to uncorrected probes was also confirmed in biological tissue by imaging neurons expressing the green fluorescence protein (GFP) in fixed brain slices (Supplementary Fig. 5).

**Fig. 4.**
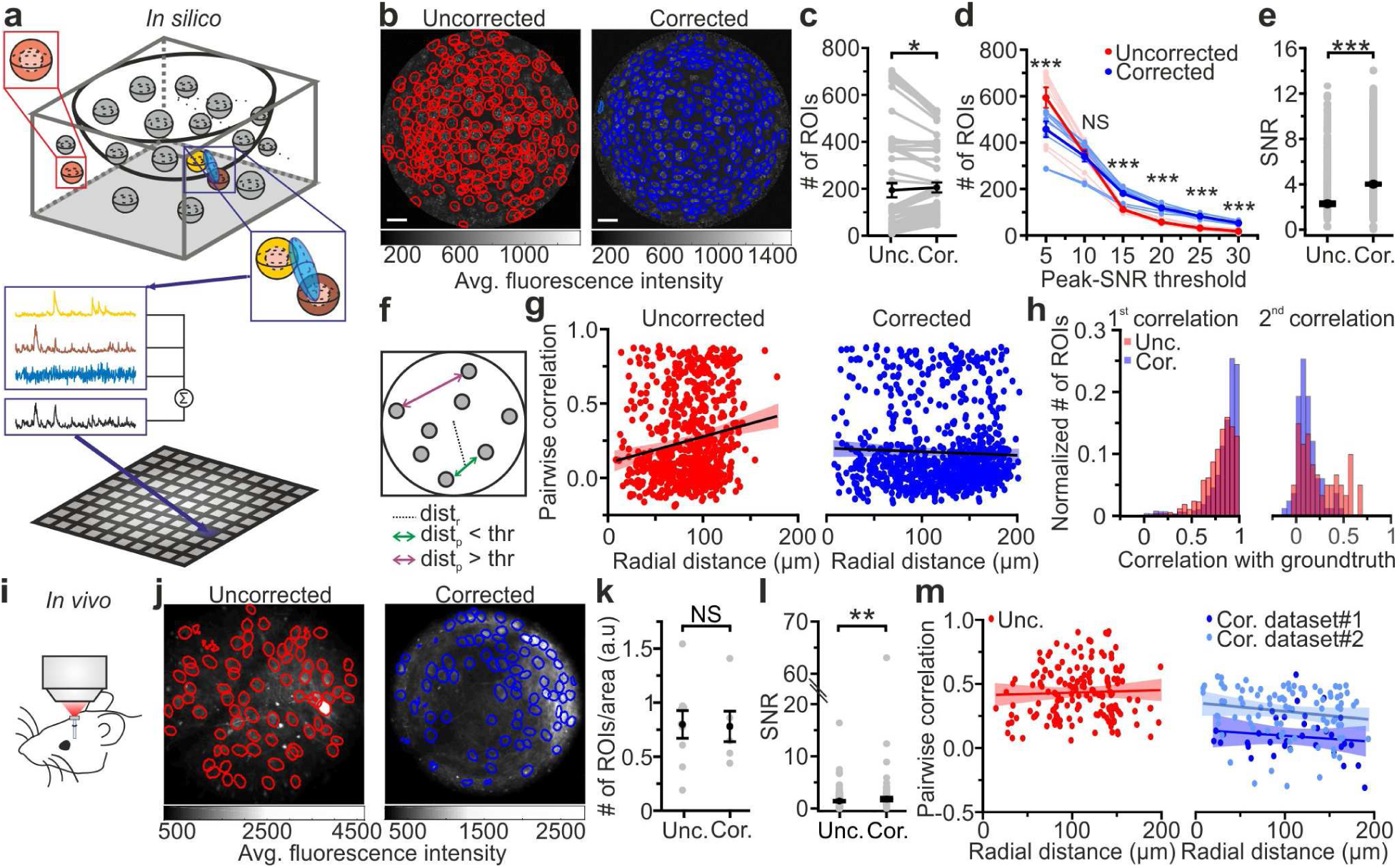
*eFOV*-microendoscopes allow higher SNR and more accurate evaluation of pairwise correlation. **a**, Schematic of the procedure for *in silico* simulation of imaging data. Neuronal activity was simulated within spheres located in a 3D volume, integrated over an elliptical PSF (blue) that was scanned on a curved FOV, projected on a 2D plane to generate artificial frames, and modulated through an intensity mask. Only voxels falling within the PSF contributed to the pixel signal (black trace). **b**, Segmentation of *in silico* data for uncorrected (left, red lines indicate identified ROIs) and corrected (right, blue lines indicate identified ROIs) endoscopes. **c**, Number of ROIs segmented in simulated FOVs for uncorrected (Unc.) and corrected (Cor.) microendoscopes. n = 54 segmentations from 9 simulated FOVs, Wilcoxon signed-rank test, p = 0.037. In this as well as in other figures, values from individual experiments are shown in grey, the average of all experiments in black and error bars indicate sem, unless otherwise stated. In this as well as in other figures: *, p < 0.05; **, p < 0.01; ***, p < 0.001; NS, not significant. **d**, Number of segmented ROIs as a function of the peak-SNR threshold in artificial data from n = 9 simulated experiments. A two-way ANOVA with interactions showed a significant effect of peak-SNR threshold (p = 1E-50) and of the interaction between peak-SNR threshold and probe type (p = 1E-5) on the number of segmented ROIs, while the effect of probe type was not significant (p = 0.096). **e**, Average SNR of calcium signals under the different experimental conditions (peak-SNR threshold = 20 for the automatic segmentation). n = 987 and 1603 ROIs for 9 simulated experiments with uncorrected and corrected microendoscopes, respectively. Mann-Whitney test, p = 5E-52. **f**, Schematic representation of how radial distance (dist_r_) and pairwise distance (dist_p_) were calculated. **g**, Pairwise correlation as a function of radial distance for simulated experiments with uncorrected (left) and corrected (right) microendoscopes. In this as well as other figures, lines show the linear regression of data. Shaded areas represent 95% confidence intervals. n = 738 and n = 869 pairwise correlations for uncorrected and corrected microendoscopes, respectively, from the n = 9 simulated experiments shown in (E). Linear regression fit of data: slope = 0.002, Student’s *t*-test, p = 2E-6 and slope = −2E-4, Student’s *t*-test, p = 0.21 for uncorrected and corrected microendoscopes, respectively. **h**, Distribution of the correlation of calcium signals with the first (left) or second (right) component of the ground truth for experiments with uncorrected and corrected microendoscopes. First component: n = 987 and 1603 ROIs for uncorrected and corrected microendoscopes, respectively, from n = 9 simulated experiments. Second component: n = 62 and 85 ROIs for uncorrected and corrected microendoscopes, respectively, from n = 9 experiments. **i**, Schematic of the experimental configuration in awake animals. **j**, Two-photon images of VPM neurons expressing GCaMP7f showing manually identified ROIs for uncorrected (left, red lines) and corrected (right, blue lines) type II microendoscopes. **k**, Spatial density of ROIs identified in *in vivo* experiments. n = 9 FOVs and 6 FOVs from 3 animals with uncorrected and corrected microendoscopes, respectively. Student’s *t*-test, p = 0.92. **l**, SNR of segmented ROIs in *in vivo* recordings. n = 557 ROIs from 9 FOVs for uncorrected microendoscopes; n = 306 from 6 FOVs for corrected microendoscopes. Mann-Whitney test, p = 0.0011. **m**, Pairwise correlation as a function of radial distance for *in vivo* experiments. Number of pairwise correlations: n = 168 from 9 FOVs, n = 36 from 6 FOVs, and n = 92 from 24 FOVs for uncorrected, corrected (dataset 1), and corrected (dataset 2), respectively. Dataset 2 was obtained from experimental sessions performed during behavioral state monitoring as in Figure 6. Linear regression fit of data: slope = 0.0002, Student’s t-test p = 0.61 for uncorrected; slope = −0.0005, p = 0.42 for dataset 1; slope = −0.0007, Student’s *t*-test p = 0.1 for dataset 2. Student’s *t*-test for comparison between uncorrected and corrected (dataset 1).

**Fig. 5.**
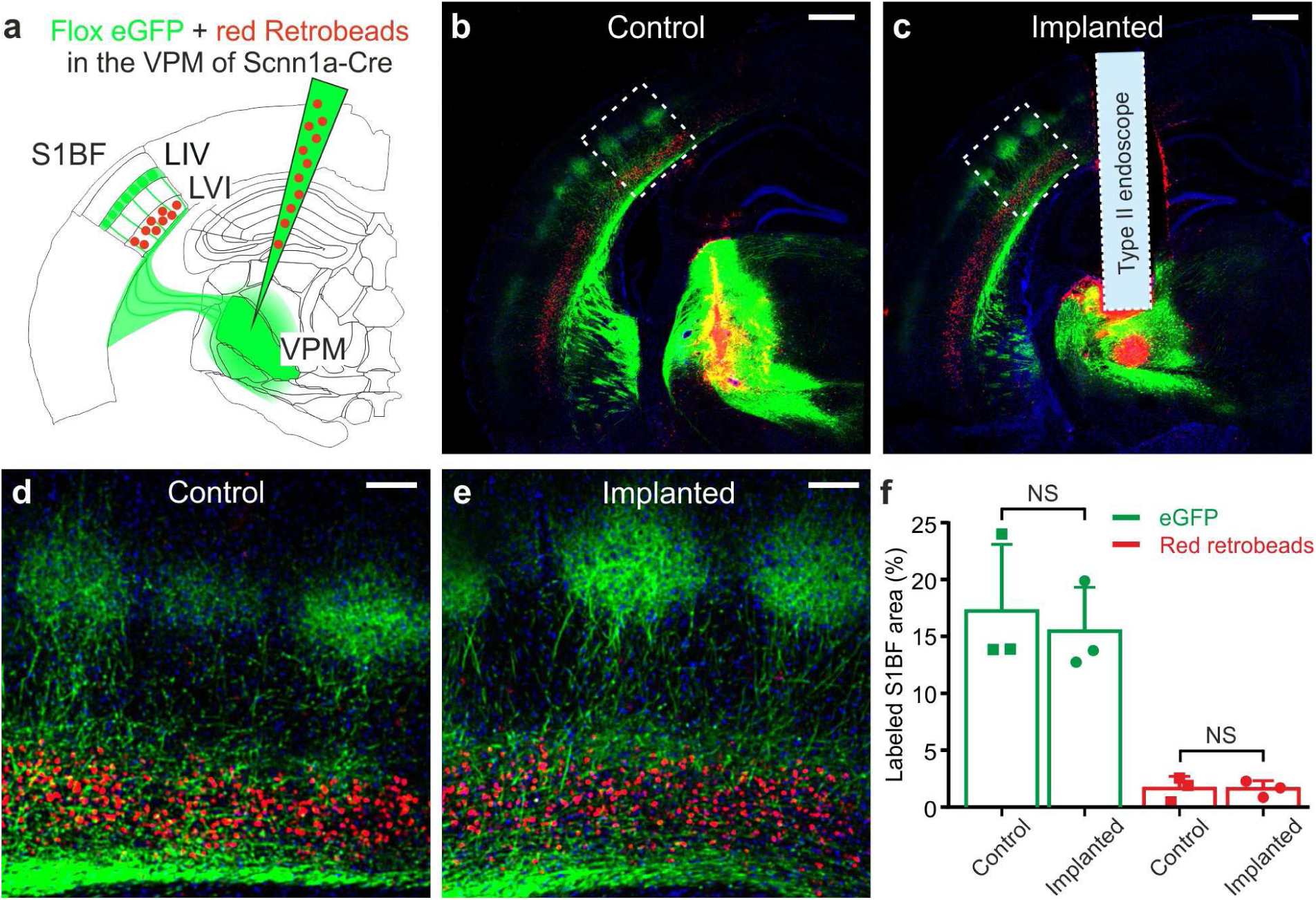
Ultrathin microendoscope implantation preserves thalamo-cortical and cortico-thalamic connectivity between S1bf and VPM. **a**, Local injection of red retrobeads and AAVs transducing floxed eGFP was performed in the VPM of Scnn1a-Cre mice. **b**, Confocal image showing a coronal slice from an injected control animal. Scale bar: 500 μm. **c**, Same as **b** but for a mouse implanted with a type II *eFOV*-microendoscope (probe diameter: 500 μm). **d**,**e**, Zoom in of the S1bf region highlighted in **b**,**c**. Scale bar: 100 μm. **f**, Percentage of labeled S1bf area with eGFP (green) and retrobeads (red) in control and implanted mice. Points indicate the value of fluorescence from 3 mice (counted 3 coronal slices from each animal), column bars indicate average ± sd. One-tailed Mann-Whitney, p = 0.20 for eGFP and p = 0.50 for red retrobeads, respectively.

### Validation of eFOV-microendoscopes for functional imaging in subcortical regions

To validate *eFOV*-microendoscopes performance for functional measurements *in vivo*, we first expressed the genetically-encoded calcium indicator GCaMP6s in the mouse hippocampal region in anesthetized mice (Supplementary Figure 6) and in the ventral posteromedial nucleus of the thalamus (VPM), a primary sensory thalamic nucleus, in awake head-restrained mice (Supplementary Figure 7). To this aim, we injected either adenoassociated viruses (AAVs) carrying a flex GCaMP6s construct together with AAVs carrying a Cre-recombinase construct under the CamKII promoter in the hippocampus of wild type mice (Supplementary Figure 6a-a_2_) or AAV carrying a floxed GCaMP6s in the VPM (Supplementary Fig. 7a,b) of Scnn1a-Cre mice ^39^, a mouse line which expresses Cre in a large subpopulation of VPM neurons. These experimental protocols resulted in the labelling of CA1 hippocampal and VPM neurons, respectively (Supplementary Figs 6,7). We imaged spontaneous activities in the CA1 hippocampal region with type I *eFOV*-microendoscopes (Supplementary Fig. 6b-d) and type III *eFOV*-microendoscopes (Supplementary Fig. 6e-g) or spontaneous activities in the VPM with the longer type II *eFOV*-microendoscopes (Supplementary Fig. 7b-d). Tens to hundreds of active ROIs *per* single FOV were identified and could be imaged using *eFOV*-microendoscopes (Supplementary Figs. 6,7).

**Fig. 6.**
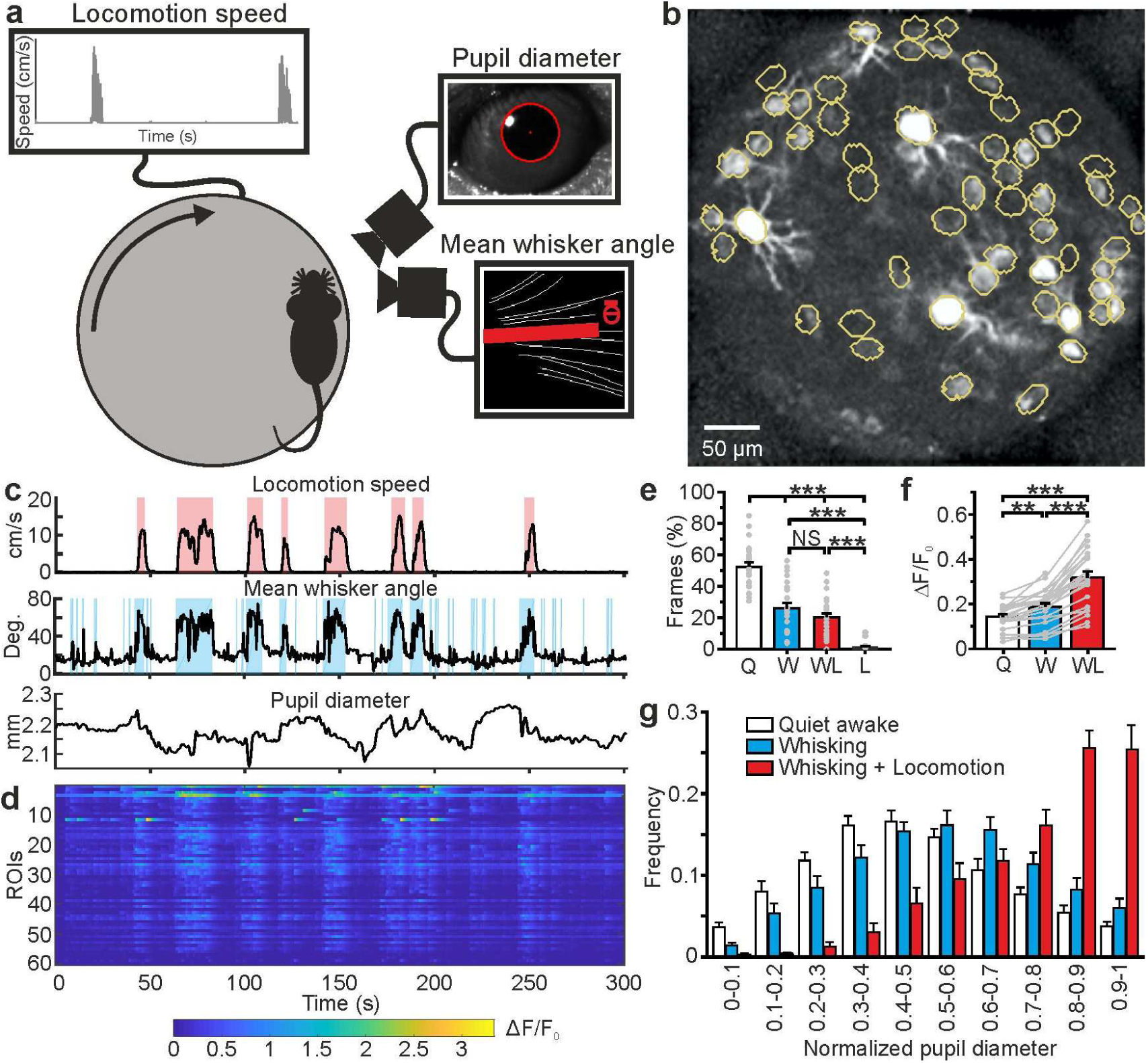
High-resolution population dynamics in the VPM of awake mice during locomotion and free whisking. **a**, Schematic of the experimental set-up for the recording of locomotion, whisker mean angle, and pupil size in awake head-fixed mice during VPM imaging using type II *eFOV*-microendoscopes. **b**, Two-photon image of GCaMP6s labelled VPM neurons *in vivo*. **c**, Representative traces of locomotion (top), whisker mean angle (middle), and pupil diameter (bottom). Red and blue shades indicate periods of locomotion and whishing, respectively (see Methods for definition). **d**, ΔF/F_0_ over time for all different ROIs in the experiment shown in **c. e**, Percentage of frames spent by the animal in quite wakefulness (Q), whisking (W), locomotion (L), and whisking + locomotion (WL). n = 24 time series from 4 animals. One-way ANOVA with Bonferroni *post hoc* correction, p = 3E-23. **f**, Average ΔF/F_0_ across ROIs under the different experimental conditions. n = 24 time series from 4 animals. One-way ANOVA with Bonferroni *post hoc* correction, p = 2E-16. **g**, Distribution of the Q, W, and WL states as a function of pupil diameter. Kolmogorov-Smirnov test for comparison of distributions of Q, W and WL states: p = 0.07 for Q *vs* W states, p = 2E-10 for Q vs WL states, p =1E-5 for W *vs* WL states. For the statistical comparison of Q, W and WL states in each range of pupil diameter a two-way ANOVA with Tukey-Kramer *post hoc* correction was performed; see Supplementary Table 3.

**Fig. 7.**
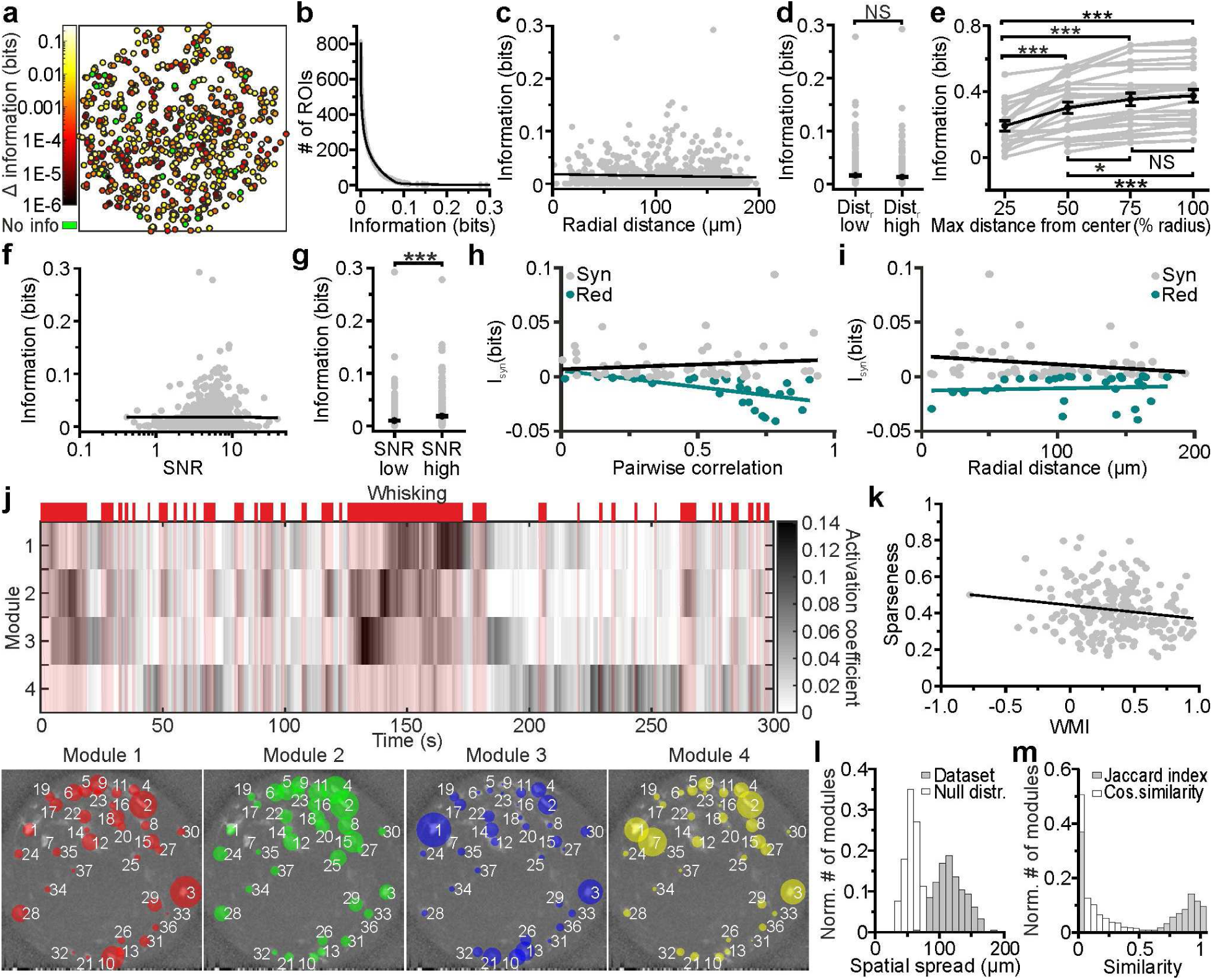
Cell-specific encoding of behaviorally-dependent information in distributed VPM subnetworks. **a**, Spatial map of neurons encoding whisking information. The pseudocolor scale shows significantly informative neurons (see Methods). Data are pooled from 24 t-series from 4 animals. See also Supplementary Fig. 8. **b**, Distribution of information content in individual ROIs. The individual ROIs information content distribution was fitted using a double exponential function (R2 = 0.99). n = 808 ROIs from 24 time series. **c**, Information content of individual neurons *vs* their radial distance from the center of the FOV. n = 842 neurons from 24 time series. Linear regression fit: slope = −3E-5, Student’s *t*-test p = 0.13. Pearson correlation coefficient: −0.053, Student’s *t*-test p = 0.13. **d**, Information content for ROIs located at low radial distance (dist_r_ low) and in lateral portions of the FOV (dist_r_ high). n = 425 and 417 ROIs from 24 t-series for dist_r_ low and dist_r_ high, respectively. Mann-Whitney test, p = 0.42. **e**, Information content of neural populations as a function of the distance from the FOV center (% radius) used to include ROIs in the population. One-way ANOVA repeated measurements with Bonferroni *post hoc*, p = 1E-18. Data pooled from 24 time series. **f**, Information content of individual neurons *vs* SNR. n = 842 neurons from 24 time series. Linear regression fit: slope = 9E-4, Student’s *t*-test p = 2.7E-4. Pearson correlation coefficient: 0.125, p = 2.7E-4. **g**, Information content for ROIs with low SNR (SNR low) and high SNR (SNR high). n = 421 and 421 ROIs from 24 t-series for dist_r_ low and dist_r_ high, respectively. Mann-Whitney test, p = 5E-5. **h**,**i**, Synergistic (grey) or redundant (dark green) information within pairs of neurons is shown as a function of pairwise correlation (**h)** and as a function of radial distance (**i**). n = 61 and 31 pairs of neurons for synergistic and redundant information content, respectively. Data from 24 time series. Linear regression fit in **h**: slope = 0.008, Student’s *t*-test p = 0.29 and slope = −0.02, Student’s *t*-test p = 6E-4 for synergistic and redundant information, respectively. Linear regression fit in **i**: slope = −0.0001, Student’s *t*-test p = 0.06 and slope = 2E-5, Student’s *t*-test p = 0.66 for synergistic and redundant information, respectively. **j**, Top: representative grayscale matrix showing the activation coefficient across time for 4 NMF modules emerging in a FOV containing 37 neurons. Periods of whisking are shown in red bars and shades. Bottom: ROIs belonging to the 4 different modules shown in the top panel. Each colored circle represents a ROI belonging to the specified module and its radius is proportional to the ROI weight within that module. The corresponding activation coefficients are presented in the upper panel. **k**, Module sparseness as a function of the whisking modulation index (WMI). n = 213 modules from 24 time series. Pearson correlation coefficient: −0.164, p = 0.016. **l**, Normalized number of modules as a function of their spatial spread for the experimental data (grey) and for a null distribution obtained by randomly shuffling the spatial position of ROIs in the FOV (white). Number of modules: n = 213 and n = 2400 for the 24 t-series of the *in vivo* dataset and for the null distribution, respectively. **m**, Normalized number of modules as a function of their Jaccard index (grey) and Cosine Similarity coefficients (white) for pairs of modules identified within the same FOV. n = 2704 modules from 24 t-series.

### Higher SNR and more precise evaluation of pairwise correlation in eFOV-microendoscopes

To establish a quantitative relationship between the improved optical properties of *eFOV*-microendoscopes and their potential advantages for precisely detecting neuronal activity, we generated two-photon imaging t-series using synthetic GCaMP data. This approach allowed us to compare results of the simulation of calcium data for both uncorrected and corrected endoscopic probes with the known ground truth of neuronal activity. Simulated neuronal activity within a volumetric distribution of cells was generated according to known anatomical and functional parameters of the imaged region (the VPM in this case) and established biophysical properties of the indicator ^40, 41^(see STAR Methods). t-series were generated by sampling simulated neuronal activity across an imaging focal surface resembling the experimental data obtained using the representative case of type II GRIN lenses for both *eFOV*-microendoscopes and uncorrected probes (Fig. 4a,b). To scan the imaging focal surface, we used an excitation volume which resembled the aberrated and aberration-corrected PSFs experimentally measured for uncorrected and *eFOV*-microendoscopes, respectively (Fig. 3, see Methods). Fluorescence traces were extracted from artificial t-series and compared between the uncorrected and corrected case. On average, a slightly larger number of ROIs could be identified in corrected probes (Fig. 4c). Crucially, we observed a nonlinear interaction between the type of probe, the number of detected ROIs, and the SNR of the calcium traces (Fig. 4d). Using corrected probes did not always allow the identification of a higher number of ROIs (p = 0.096, two-way ANOVA with respect to probe type). Rather, the use of corrected probes allowed to segment more ROIs with high SNR, shifting the distribution of SNR across ROIs to higher mean SNR values (Fig. 4e, p = 1E-5 for ANOVA with respect to the interaction).

Pairwise correlation between nearby neurons (distance between the center of neurons < 20 µm) should not vary with the radial distance because in our simulations this value was constant across neurons. However, we found an artefactual increase of correlation strength with the radial distance of neuronal pairs in uncorrected endoscopes due to the cross-contamination of activity at different points generated by the larger and aberrated PSF without the corrective lens. In contrast, correlation strength remained constant in *eFOV*-microendoscopes (Fig. 4f,g), suggesting that the PSF of the corrected probes was small enough across the FOV to decrease contamination of activity across neurons. This result is thus in agreement with the decreased spatial resolution observed in more distal parts of the FOV in uncorrected probes and the improved and homogeneous resolution across the FOV that is instead found in corrected microendoscopes (Fig. 3). Overall, these findings suggest that signal corresponding to individual neurons could be more accurately extracted from ROIs across the FOV in corrected microendoscopes. We quantified this in synthetic data by evaluating, for each ROI, the correlation of the extracted calcium trace with the ground truth fluorescence signal generated by the simulated neuronal activity contained in that ROI (Fig. 4h left). For those ROIs whose fluorescence dynamics were determined by more than one neuron, the correlation with the second most relevant cell was also calculated (Fig. 4 h right). We found that ROIs segmented from *eFOV*-microendoscopes displayed larger correlation with the ground truth signal of an individual neuron (Fig. 4h left) and lower correlation with the ground truth signal of other nearby neurons (Fig. 4h right) compared to uncorrected probes, suggesting that aberration correction allowed to collect more precisely from single cellular emitters and decreased cross-contamination between neurons.

We experimentally validated the results of the simulations performing functional imaging in the VPM using type II *eFOV*-microendoscopes before and after the removal of the corrective microlens in awake head-restrained mice (Fig. 4i,j). The number of ROIs detected under the two conditions was not significantly different (Fig. 4k). However, we found increased average SNR of calcium signals in corrected compared to uncorrected probes (Fig. 4l), confirming also in experimental data (as it happens in simulations, Fig, 4d,e) that the use of an *eFOV*-microendoscope shifts the distribution of SNR across ROIs towards having a higher proportion of ROIs with large SNR value. Moreover, pairwise correlation increased as a function of the radial distance of the pair in uncorrected probes compared to corrected ones (Fig. 4m), in agreement with the analysis of the artificial calcium t-series (Fig.4g). Overall, results of simulations and experiments demonstrate that correcting optical aberrations in *eFOV*-microendoscopes enabled higher SNR and more precise evaluation of pairwise correlation compared to uncorrected probes.

### Spatial mapping of behavior state-dependent information in sensory thalamic nuclei in awake mice

We then focused our attention on the VPM, a key region which relays somatosensory (whisker) information to the barrel field of the primary somatosensory cortex (S1bf) through excitatory thalamocortical fibers ^42^. VPM also receives strong cortical feedback from corticothalamic axons of deep cortical layers. Cortical inputs to VPM has been proposed to strongly modulate thalamic activity. Thus, to study VPM physiology it is fundamental to preserve corticothalamic and thalamocortical connectivity. Electrophysiological recordings showed that VPM networks are modulated by whisking and behavioral state ^43-45^. However, how information about whisking and other behavioral state-dependent processes (e.g arousal, locomotion) are spatially mapped in VPM circuits at the cellular level is largely unknown. We used *eFOV*-microendoscopes to address this question.

As an important control experiment, we first confirmed that the ultrathin GRIN lenses that we used in our study (diameter ≤ 500 µm) did not significantly damage anatomical thalamocortical and corticothalamic connectivity, a difficult task to achieve with larger cross-section GRIN lenses or with chronic optical windows (Supplementary Fig. 7e). To this aim, we performed local co-injections in the VPM of Scnn1a-Cre mice of red retrobeads to stain corticothalamic projecting neurons with axons targeting the VPM and of an adenoassociated virus carrying a floxed GFP construct to stain thalamocortical fibers (Supplementary Fig. 5a). We evaluated the amount of thalamocortical and corticothalamic connectivity looking at the percentage of pixels displaying green and red signal in the S1bf region in endoscope-implanted *vs* non-implanted mice (Fig. 5a-c). In accordance with the known anatomy of the thalamocortical system ^46^, we found that the green signal was mostly localized in layer IV barrels and in layer V/VI while the red signal was largely restricted to layer VI (Fig. 5d,e). Importantly, we found no difference in the percentage of pixels displaying green and red signals in implanted *vs* non-implanted mice (Fig. 5f).

We then used *eFOV*-microendoscopes to address the question of how information about motor behavior (e.g., locomotion and whisking) and internal states (e.g., arousal state) are mapped on VPM circuits at the cellular level. To this aim, we used *eFOV*-microendoscopes to perform GCaMP6s imaging in VPM circuits in awake head-restrained mice while monitoring locomotion, whisker mean angle, and pupil diameter (Fig. 6a-d, see Methods). We identified quiet (Q) periods, time intervals that were characterized by the absence of locomotion and whisker movements, and active (A) periods, intervals with locomotor activity, dilated pupils, and whisker movements. Active periods were further subdivided into whisking (W), whisking and locomotion (WL), and locomotion with no whisking (L). Fig.6e shows a histogram representing the amount of time spent in the different behavioral states. Mice whisk when they move, therefore L periods were rare. We found that Q periods showed calcium events that were sparsely distributed both across time and neurons (Fig. 6c,d). In contrast, active periods displayed an increase in both frequency and amplitude of calcium signals across VPM neurons compared to Q periods (average frequency: f_Q_ = 1.95 ± 0.02 Hz, f_A_ = 2.22 ± 0.02 Hz, Student’s *t*-test p = 2E-74, n = 24 t-series from 4 mice; average amplitude: A_Q_ = 0.137 ±0.005 ΔF/F_0_, A_A_ = 0.245 ± 0.008 ΔF/F_0_, Student’s *t* -test p = 2E-128, n = 24 t-series from 4 mice). This resulted in a significant increase in the average fluorescence across neurons during the active W and WL periods compared to Q periods (Fig. 6f). The increase in the frequency of WL also correlated with pupil size (Fig. 6g and Supplementary Table 3), which reflects the arousal level of the animal ^47^.

### Cell-specific encoding of whisking-dependent information in distributed VPM subnetworks

We investigated how neuronal activity was modulated by an important behavioral variable: whether the mouse was whisking or not. We considered neuronal activity both at the single-cell and population level. We quantified the content of mutual information about whisking state (whether or not the mouse was whisking; shortened to whisking information hereafter) based on the fluorescence signals extracted from individual neurons (Fig. 7a). We found that many neurons were informative about whisking, but only a fraction were particularly informative (Fig. 7b). Highly-informative neurons were sparse and could be surrounded by low-information-containing neurons (Fig. 7a). This indicates that while informative neurons are distributed across the FOV, information is strongly localized and highly cell-specific. Whisking information in individual cells measured with the *eFOV*-microendoscope was constant over the radial distance (Fig. 7b,c). Moreover, adding neurons recorded at higher radial distances to a population decoder improved the amount of extracted whisking information compared to considering only neurons in the center of the FOV (Fig. 7e). Together, these results suggest that the corrected endoscope was effective at capturing information also at the FOV borders. Information of single cells correlated positively with the SNR of their calcium signals (Fig. 7f), with higher amount of information carried, on average, by cells displaying higher SNR (Fig. 7c). This suggests that the benefit of *eFOV*-microendoscopes in providing higher SNR, demonstrated earlier, also translates in an ability to extract more information about the circuit’s encoding capabilities (Fig. 7e).

We also considered the redundancy and synergy of whisking information carried by pairs of simultaneously recorded nearby (distance between neurons < 20 µm) neurons. This analysis is important because how pairwise correlations shape the redundancy and synergy of information representation is fundamental to the understanding of population codes ^10, 11, 48^. We found that redundancy and synergy of pairs recorded with the *eFOV*-microendoscopes were, on average, smaller and larger in pairs with stronger correlations, respectively (Fig. 7h). Importantly, and as a consequence of the fact that *eFOV*-microendoscopes avoid the artificial increase of pairwise correlations close to the FOV borders, we did not observe an increase of synergy or redundancy between nearby neurons close to the FOV border (Fig. 7i). This shows that aberration correction helps avoiding the generation of artificially biased estimates of synergy and redundancy near the FOV border.

We finally turned to analyzing the properties of firing at the level of the whole population recorded in the FOV. We applied non-negative matrix factorization (NMF)^49^ to identify subpopulations of neurons (modules, Fig. 7j) characterized by correlated activity. Detected modules were differentially activated in time and were sparsely distributed in space (Fig. 7j). Moreover, we found that modules could be oppositely modulated by whisking, with the activity of some modules being enhanced and the activity of some other modules being depressed by whisking (Fig. 7j). We computed the whisking modulation index (WMI, see STAR Methods for definition) and found that the large majority of modules was positively modulated by whisking (WMI > 0 for 89.6 ± 0.4 % of total modules number, n = 24 FOVs), while the activity of a minority of modules was suppressed during whisking periods (negatively modulated, WMI < 0 for 10.4 ± 0.4 % of total modules number, n = 24 FOVs). Sparseness of modules appeared to be negatively correlated with the WMI, suggesting that those few modules that were negatively modulated by whisking were also characterized by few, but highly informative neurons (within the ensemble) (Fig. 7k). In contrast, modules with high WMI values were less sparse, suggesting similar activity (and information) across most of the neurons belonging to these ensembles. Single modules covered distances of hundreds of μm spanning the whole *eFOV* (Fig. 7l). We showed that the spatial distances covered by functional modules were higher than distances obtained by chance, using a permutation test (Fig. 7l). This suggests that corrected probes allow unveiling functional relationship between groups of neurons spanning the whole *eFOV*. Neurons could belong to different ensembles as quantified by the distribution of the values of the Jaccard index among pairs of modules (Fig. 7m). This distribution was bimodal, with a peak at zero (no ROI belonged to both modules) and the other peak towards the value 1 (the two modules were composed by the same ROIs). However, when ROIs belonged to more than one module they tended to have module-specific weight (i.e., different weights for different modules). In fact, the distribution of values for the cosine similarity, an index which considers the weight of ROIs within a module (see Methods), was shifted toward smaller values compared to the distribution of the Jaccard index values (Fig. 7m).

## Discussion

In this study, we designed, developed, characterized, and successfully validated a new approach to correct aberrations in ultrathin GRIN-based endoscopes using aspheric lenses microfabricated with 3D micro-printing based on TPL ^33^. We applied aberration-corrected microendoscopes to study how free whisking and behavior state-dependent information is encoded in primary sensory thalamic nuclei in awake mice. We reported the spatial map of encoding of such information in the VPM nucleus, maintaining unaltered the bidirectional cortico-thalamic connectivity.

GRIN lenses, alone or in combination with fiber-bundles, have been used to perform one- and two-photon endoscopic imaging in deep brain areas, such as the hippocampus ^50-52^, the striatum ^27^, and the hypothalamus ^27, 53, 54^. GRIN microendoscopes have also been used to perform simultaneous functional imaging of two different brain regions ^55^, allowing concurrent monitoring of neuronal dynamics in areas otherwise not accessible with single FOV systems. Even though the combination of GRIN lenses with two-photon imaging allows improved optical sectioning, most GRIN endoscopes have been operating with limited performance due to optical aberrations. These aberrations derive from the intrinsic nonaplanatic properties of GRIN rods ^56^ and limit the spatial resolution and the usable FOV ^25, 28, 30, 32^. Aberrations typically increase with the length and NA of the GRIN lens, making high-resolution imaging of large FOV in deeper areas a challenging goal. Aberrations can be partially compensated by adding additional optical elements, such as a cover glass ^57^, a single high refractive index planoconvex lens ^25^, and multiple planoconvex lenses combined with diffractive optical elements ^32^. By adding a single planoconvex ball lens to the distal end of a customized GRIN rod (rod diameter: 1 mm), Barretto et al. (2009) increased the NA up to 0.82 correcting on-axis aberrations and they imaged neuronal dendritic spines in GFP-expressing hippocampal pyramidal neurons in live mice ^25^. To correct off-axis aberrations while maintaining high NA, at least a second planoconvex lens was needed ^32^. Fabrication of high performance multi-element optical systems for on-axis and off-axis aberration correction, however, requires high precision in the fabrication and assembly of the corrective optical elements. Moreover, it usually requires the use of external metal cannulas to hold the various optical elements aligned and provide mechanical stability to the optical system. For this reason, so far, aberration correction with built-in optical elements was limited to GRIN lenses of diameter ≥ 1 mm and overall endoscopic probe diameter (GRIN + cannula) of ≥ 1.4 mm ^25, 32^. Improving optical performances in ultrathin (diameter ≤ 0.5 mm) microendoscopes with built-in optical elements thus remains a major technical challenge. Since the insertion of the probe irreversibly damages the tissue above the target area, reducing the size of the probe and consequently its invasiveness is of utmost importance when imaging deep brain regions.

We devised a new approach to solve this problem and used 3D micro-printing based on TPL ^33, 58^ to microfabricate polymeric aspheric lenses that effectively corrected aberrations in ultrathin GRIN-based endoscopes. Corrective lenses were first fabricated on glass coverslips, which were aligned and assembled with the GRIN rod to form an aberration-corrected microendoscope. This optical design resulted in improved axial resolution and extended effective FOV (Fig. 3), in good agreement with the predictions of the optical simulations (Fig. 2). Aberration correction in GRIN microendoscopes can be achieved using adaptive optics (AO) ^28, 30, 31, 56^. For example, using pupil-segmentation methods for AO, diffraction-limited performance across an enlarged FOV was obtained in GRIN-based endoscopes with diameter of 1.4 mm ^28, 30^ and, in principle, this approach could be extended to probes with smaller diameter. Nevertheless, AO through pupil segmentation requires significant modification of the optical setup and the use of an active wavefront modulation system (e.g. deformable mirror device or liquid crystal spatial light modulator) which needs the development of ad-hoc software control. Moreover, AO through pupil segmentation may limit the temporal resolution of the system, since multiple AO corrective patterns must be applied to obtain an aberration-corrected extended FOV ^28^. Compared to AO approaches, the technique developed in this study does not necessitate modification of the optical path nor the development of ad-hoc computational approaches. Moreover, it is easily coupled to standard two-photon set-ups, and does not introduce limitations in the temporal resolution of the imaging system.

Using synthetic calcium data, we demonstrated that the improved optical properties of *eFOV*-microendoscopes directly translate into important advantages for measuring neural population parameters *in vivo*. Namely, they achieve a higher SNR of calcium signals and a more precise evaluation of pairwise correlation compared to uncorrected GRIN lenses, predictions that were all confirmed experimentally in awake mice (Fig. 4). Importantly, synthetic calcium data also allowed us to evaluate the impact of correcting optical aberration on the accuracy in extracting neuronal activity and population codes from calcium imaging data. We found larger correlation of extracted calcium traces with the known ground truth of neuronal spiking activity in *eFOV*-microendoscopes compared to uncorrected probes. Traces extracted from *eFOV*-microendoscopes were less contaminated by neighboring neurons compared to uncorrected probes, in agreement with the higher and more homogeneous spatial resolution of *eFOV*-microendoscopes (Fig. 4). All these achievements were obtained without increasing the lateral size of the probe, thus minimizing tissue damage in biological applications.

Studying neuronal population codes requires the measurement of neuronal population activity with high precision, large SNR, and without introducing artificial bias on the activity of individual neurons and the measures of relationship between them, such as pairwise correlations. In particular, pairwise correlations are thought to be fundamental for information coding, signal propagation, and behavior ^1-3, 5, 10^. Here, we demonstrate that the homogeneous spatial resolution, which characterized *eFOV*-microendoscopes (Fig. 3), allowed an unbiased computation of pairwise correlations, a higher correlation of extracted calcium traces with the ground truth neuronal activity, and a smaller contamination of extracted signals by neighboring cells (Fig. 4).Several studies suggested that even small biases in measuring single cell and pairwise properties, either, for example, in incorrectly measuring the average amount of correlations or the heterogeneity of single cell tuning and of correlation values, may lead to large biases in determining how these populations encode information ^4, 5, 59^. *eFOV*-microendoscopes allow to remove the artefacts and biases introduced by uncorrected GRIN endoscopes in measuring both individual cell information properties and correlation between each and every pair of neurons. The advantages introduced by *eFOV*-microendoscopes are therefore essential for unraveling the true nature of population codes in deep brain structure.

We used the unique features of the *eFOV*-microendoscopes to study how highly correlated activity is mapped in the VPM thalamic nucleus of awake mice with unprecedented combination of high spatial resolution across the FOV and minimal invasiveness (Figs. 5-7). The VPM is a primary somatosensory thalamic nucleus which relays sensory information to the S1bf ^60^. However, VPM also receives strong cortical innervation which deeply affects VPM activity ^61-63^. We first showed that the small cross-section of the *eFOV*-microendoscopes developed here preserved thalamocortical and corticothalamic connectivity (Fig. 5), a fundamental prerequisite for VPM physiology and a hardly achievable task with larger cross section GRIN lenses or with chronic windows (Supplementary Fig. 7e). We then imaged GCaMP6s-expressing VPM neurons while monitoring locomotion, whisker movement, and pupil diameter (Fig. 6). We found cell-specific encoding of whisking information in distributed functional VPM subnetworks. Most individual neurons encoded significant amount of whisking information generating distributed networks of informative neurons in the VPM (Fig. 7). However, the amount of encoded information was highly cell-specific, with high-information-containing neurons being sparsely distributed in space and surrounded by low-information-containing cells. Sparse distribution of information content has been similarly observed in other brain areas ^10, 64^. At the population level, we observed that whisking modulated functional ensembles of neurons, which were oppositely modulated by whisking. Some ensembles displayed enhanced activity upon whisking, while some other ensembles showed suppressed activity by whisking. Interestingly, single neurons could belong to multiple functional ensembles, but their weight within one ensemble was ensemble-specific (Fig. 7). Overall, the application of *eFOV*-microendoscopes revealed for the first time the complexity and cellular specificity of the encoding of correlated behavioral state-dependent information in a primary thalamic sensory nucleus.

In summary, we developed a new methodology to correct for aberrations in ultrathin microendoscopes using miniutarized aspheric lenses fabricated with 3D printing based on TPL. This method is flexible and can be applied to the GRIN rods of different diameters and lengths that are required to access the numerous deep regions of the mammalian brain. Corrected endoscopes showed improved axial resolution and up to 9 folds extended effective FOV, allowing high resolution population imaging with minimal invasiveness. Importantly, we demonstrate that *eFOV*-microendoscopes enable more precise extraction of population codes from two-photon imaging recordings. Although *eFOV-*microendoscopes have been primarily applied for functional imaging in this study, we expect that their use can be extended to other applications. For example, *eFOV-* microendoscopes could be combined with optical systems for two-photon patterned optogenetic manipulations ^65-67^ and for simultaneous functional imaging and optogenetic perturbation ^68-72^. Moreover, besides its applications in the neuroscience field, *eFOV*-microendoscopes could be used in a large variety of optical applications requiring minimally invasive probes, ranging from cellular imaging ^34, 73^ to tissue diagnostic ^74, 75^. Importantly, applications of ultrathin *eFOV-* microendoscopes to other fields of research will be greatly facilitated by the built-in aberration correction method that we developed. This provides a unique degree of flexibility that allows using ready-to-use miniutarized endoscopic probes in a large variety of existing optical systems with no modification of the optical path.

## Materials and methods

### Animal models

Experimental procedures involving animals have been approved by the Istituto Italiano di Tecnologia Animal Health Regulatory Committee, by the National Council on Animal Care of the Italian Ministry of Health (authorization # 1134/2015-PR, # 689/2018-PR) and carried out according to the National legislation (D.Lgs. 26/2014) and to the legislation of the European Communities Council Directive (European Directive 2010/63/EU). Experiments were performed on adult (8-14 week old) mice. C57BL/6J mice (otherwise called C57, Charles River #000664, Calco, IT) were used in Supplementary Fig. 6. Data reported in Figs. 4-7 and Supplementary Figure 7 were obtained from B6;C3-Tg(Scnn1a-cre)3Aibs/J (JAX #009613, Jackson Laboratory, Bar Harbor, USA) mice crossed with C57 mice (otherwise called Scnn1a-Cre). Both male and female animals were used in this study. Animals were housed in individually ventilated cages under a 12-hr light:dark cycle. Access to food and water was *ad libitum.* The number of animals used for each experimental dataset is specified in the text or in the corresponding Figure legend.

## Methods details

### Design and simulation of corrective lenses and of *eFOV*-microendoscopes

Simulations were run with OpticStudio15 (Zemax, Kirkland, WA) to define the profile of the aspheric corrective lens to be integrated in the aberration corrected microendoscopes, with the aim to achieve: *i)* a full-width half maximum (FWHM) lateral resolution < 1 µm at the center of the FOV; *ii)* a FWHM axial resolution below < 10 µm; *iii)* a working distance between 150 µm and 220 µm into living brain tissue. The wavelength used for simulations was λ = 920 nm. The surface profile of corrective aspheric lenses was described in ^76^:

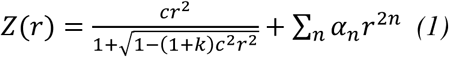

Since GRIN lenses have intrinsic spherical aberration, the optimization for the shape of the corrective lenses started with the profile of a Schmidt corrector plate ^77^ as initial guess; the parameters *c, k, α*_*n*_ (with n = 1-8) in equation *(1)* were then automatically varied in order to maximize the Strehl ratio ^78^ over the largest possible area of the FOV (Supplementary Table 1). A fine manual tuning of the parameters was performed for final optimization.

### Corrective lens manufacturing and microendoscope assembly

The optimized aspheric lens structure obtained with simulations was exported into a 3D mesh processing software (MeshLab, ISTI-CNR, Pisa, IT) and converted into a point cloud dataset fitting the lens surface (with ∼ 300 nm distance among first neighborhood points). Two-photon polymerization with a custom set-up ^33^ including a dry semi-apochromatic microscope objective (LUCPlanFLN 60x, NA 0.7, Olympus Corp., Tokyo, JP) and a near infrared pulsed laser beam (duration: 100 fs; repetition rate: 80 MHz; wavelength: 780 nm; FemtoFiber pro NIR, Toptica Photonics, Graefelfing, DE) was used for the fabrication of the corrective lenses. A drop of resin (4,4’-Bis(diethylamino)benzophenone photoinitiator mixed with a diacrylate monomer), sealed between two coverslips, was moved by a piezo-controlled stage (model P-563.3CD, PI GmbH, Karlsruhe, DE) with respect to the fixed laser beam focus, according to the 3D coordinates of the previously determined point cloud with precision of 20 nm. Output laser power was ∼ 15 mW at the sample. Once the surface was polymerized, the lens was dipped for ∼ 2 minutes in methanol followed by ∼ 1 minute immersion in isopropyl alcohol and finally exposed to UV light (λ = 365 nm; 3 Joule / cm^2^) to fully polymerize the bulk of the structure.

For fast generation of multiple lens replicas, a molding ^37^ technique was used. To this end, polydimethylsiloxane (PDMS, Sylgard 164, 10:1 A:B, Dow Corning, Auburn, MI) was casted onto the lens and hardened by heat cure in a circulating oven at 80°C for approximately 30 minutes. The resulting bulked structure of solid PDMS was then used as negative mold. A drop of a UV-curable optically-clear adhesive with low fluorescent emissivity (NOA63, Norland Products Inc., Cranbury, NJ) was deposited on the negative mold, pressured against a coverslip (diameter: 5 mm) of appropriate thickness (thickness: 100 or 200 µm depending on the *eFOV*-microendoscope type, Fig. 1) and hardened by UV exposure. The coverglass with the lens attached was detached from the mold and glued onto a metal ring. One end of the appropriate GRIN rod (NEM-050-25-10-860-S; NEM-050-43-00-810-S-1.0p; GT-IFRL-035-cus-50-NC; NEM-035-16air-10-810-S-1.0p, Grintech GmbH, Jena, DE) was attached perpendicularly to the other surface of the coverslip using NOA63. Alignment of the corrective lens and the GRIN rod was performed under visual guidance using an opto-mechanical stage, custom-built using the following components (Supplementary Fig. 2): camera (DCC1645C, Thorlabs, Newton, NJ), fine z control (SM1Z, Thorlabs, Newton, NJ), coarse z control (L200/M, Thorlabs, Newton, NJ), xyz control (MAX313D/M, Thorlabs, Newton, NJ), high power UV LED (M375L3, Thorlabs, Newton, NJ), long pass dichroic mirror (FF409-Di02, Semrock, Rochester, NY), tube lens (AC254-150-A, Thorlabs, Newton, NJ), objective (UPlanFLN 4x 0.13NA, Olympus, Milan, IT), xy control (CXY1, Thorlabs, Newton, NJ), custom GRIN rod holder, and fiber optic holder (HCS004, Thorlabs, Newton, NJ). An additional and removable coverglass or a silicone cap (Kwik-Cast Sealant, World Precision Instruments, Friedberg, DE) was glued on the top of every support ring to keep the polymeric corrective lens clean and to protect it from mechanical damage.

### Optical characterization of *eFOV*-microendoscopes

Optical characterization of *eFOV*-microendoscopes was carried out with a two-photon laser-scanning microscope equipped with a wavelength-tunable, ultrashort-pulsed, mode-locked Ti:Sapphire laser source (Ultra II Chameleon, pulse duration: 160 fs; repetition rate: 80 MHz; wavelength: 920 nm; Coherent Inc., Santa Clara, CA) and a commercial Prairie Ultima IV scanhead (Bruker Corporation, Milan, IT). For all measurements, the wavelength was set at 920 nm. The optomechanical assembly used for the *eFOV*-microendoscope characterization is shown in Supplementary Fig. 2c. The coupling objective was EC Epiplan-Neofluar 20x, 0.5NA (Zeiss, Oberkochen, DE). The z control (SM1Z) and xy control (CXY2) were purchased from Thorlabs (Newton, NJ). Spatial resolution of each microendoscope was evaluated using subresolved spherical fluorescent beads (diameter: 100 nm, Polyscience, Warrington, PA), following a previous spatial calibration using a custom fluorescent ruler (Motic, Xiamen, CN). The same ruler was used to evaluate the distortion of the FOV. To visualize the curvature of the imaging field, thin (thickness: 300 nm) fluorescent slices ^38^ were used. Fluorescent samples were deposited on a microscope slide and imaged through the endoscope assembly aligned to the microscope objective, with or without the corrective microlens above the coverslip, using the coupling apparatus described in Supplementary Fig. 2. Imaging was performed with the distal end of the GRIN rod immersed in a droplet of water placed on the slide. The distance between the focal plane of the microscope objective and the endoscope assembly was ∼ 100 µm and it was fixed for all measurements. Given the imaging field curvature of endoscopes, for both the ruler and the thin fluorescent slices (planar samples), the acquisition of z-series of images (512 pixels x 512 pixels, with 1 µm axial step) was performed.

### Viral injections and microendoscope implantation

Adeno-associated viruses (AAVs) AAV1.Syn.flex.GCaMP6s.WPRE.SV40, AAV1.CAG.Flex.eGFP.WPRE.bGH, AAV1.CaMKII0.4.Cre.SV40 were purchased from the University of Pennsylvania Viral Vector Core. AAV1.Syn.flex.GCaMP7f.WPRE.SV40 was purchased from Addgene (Teddington, UK) Animals were anesthetized with isoflurane (2 % in 1L/min O_2_), placed into a stereotaxic apparatus (Stoelting Co, Wood Dale, IL) and maintained on a warm platform at 37°C. The depth of anesthesia was assessed by monitoring respiration rate, heartbeat, eyelid reflex, vibrissae movements, reactions to tail and toe pinching. 2% lidocaine solution was injected under the skin before surgical incision. A small hole was drilled through the skull and 0.5 - 1 µl (30 - 50 nl/min, UltraMicroPump UMP3, WPI, Sarasota, FL) of AAVs containing solution was injected at stereotaxic coordinates: 1.4 mm posterior to bregma (P), 1 mm lateral to the sagittal sinus (L), and 1 mm deep (D) to target the hippocampal CA1 region; 1.7 mm P, 1.6 mm L, and 3 mm D to target the VPM. Co-injection of AAV1.Syn.flex.GCaMP6s.WPRE.SV40 and AAV1.CaMKII0.4.Cre.SV40 (1:1) was performed to express GCaMP6s in hippocampus CA1 pyramidal cells of C57 mice (Supplementary Fig. 6). Injection of AAV1.Syn.flex.GCaMP6s.WPRE.SV40 (1:4 in saline solution) (Figs 6-7 and Supplementary Fig. 7) or AAV1.Syn.flex.GCaMP7f.WPRE.SV40 (1:4 in saline solution) (Fig. 4) in the Scnn1a-Cre mice was performed to express GCaMP6/7 in the VPM. Following virus injection a craniotomy (∼ 600 x 600 µm^2^ or ∼ 400 x 400 µm^2^ depending on the endoscope size) was performed over the neocortex at stereotaxic coordinates: 1.8 mm P and 1.5 mm L to image the hippocampus; 2.3 mm P and 2 mm L to reach the VPM. A thin column of tissue was suctioned with a glass cannula (ID, 300 µm and OD, 500 µm; Vitrotubs, Vitrocom Inc., Mounting Lakes, NJ) and the microendoscope was slowly inserted in the cannula track, using a custom holder, down to the depth of interest and secured by acrylic adhesive and dental cement to the skull. If necessary, metal spacers (thickness: ∼ 100 µm) were glued on the flat coverslip surface to obtain the desired protrusion distance of the GRIN rod. For experiments in awake animals (Figs 4,6,7 and Supplementary Fig. 7), a custom metal head plate was sealed on the skull using dental cement to assure stable head fixation during two-photon imaging. An intraperitoneal injection of antibiotic (BAYTRIL,Bayer, DE) and dexamethasone (MSD Animal Health, Milan, IT) was performed to prevent infection and inflammation. Animals were then positioned under a heat lamp and monitored until recovery.

To evaluate thalamocortical (TC) and corticothalamic (CT) anatomical connectivity (Fig. 5) after implantation of microendoscopes, Scnn1a-Cre mice were injected, as described above, at 100 nl/min (UltraMicroPump UMP3, WPI, Sarasota, FL) with AAV1.CAG.Flex.eGFP.WPRE.bGH virus (1:2 in saline solution) and red retrobeads (1:8 in saline solution, Fluorescent Latex Microspheres, LumaFluor Inc., Durham, NC) in VPM after tissue suctioning and in the absence of tissue suctioning. The total injected volume was 250 nl.

### Functional imaging with *eFOV*-microendoscopes *in vivo*

For experiments in anesthetized conditions (Supplementary Fig. 6), three to five weeks after injection, mice were anesthetized with urethane (16.5%, 1.65 g*kg^-1^) and placed into a stereotaxic apparatus to proceed with imaging. Body temperature was measured with a rectal probe and kept at 37°C with a heating pad. Depth of anesthesia was assured by monitoring respiration rate, eyelid reflex, vibrissae movements, and reactions to pinching the tail and toe. In some experiments, oxygen saturation was controlled by a pulseoxymeter (MouseOx, Starr Life Sciences Corp., Oakmont, PA).

For experiments in behaving mice (Figs 4,6,7 and Supplementary Fig. 7), imaging was performed two to four weeks after the endoscope implant, following 7-10 days of habituation, in which mice were placed daily on the set up, each day for a longer time duration, up to 45 minutes ^79^. Mice were allowed to run spontaneously on the wheel. During experiments, recording sessions were up to five 5 minutes long (frame rate: ∼ 3 Hz) and they were interleaved by 5 minutes in which no imaging was performed. For scanning imaging of GCaMP6/7-expressing neurons, the same microscope set-up used for the optical characterization of *eFOV*-microendoscopes was used and GCaMP6/7 fluorescence was excited at 920 nm (laser power: 28-90 mW).

### Measurement of whisker angle, pupil size, and locomotion

In Figs 6-7, whimaged spontaneous activitiesimaged spontaneous activitiesisker movements in the whisker pad contralateral to the recording site were imaged with a high-speed,Basler acA800 (Basler, Ahrensburg, DE; acquisition rate: 150 Hz) through a 45° tilted mirror placed below the whiskers. Illumination was provided by an array of infrared LEDs (emission wavelength: 800 nm) fixed to the microscope objective and aligned to the whiskers and the mirror. Imaging of the contralateral eye was performed during each experimental session with a Basler acA800 (Basler, Ahrensburg, DE), coupled with a long pass filter (HP 900, Thorlabs, Newton, NJ) at 80 Hz acquisition rate. Illumination light was provided by the pulsed laser sourced which was used to perform two-photon microendoscopic imaging (λ = 920 nm). Locomotor activity was measured with an optical encoder (AEDB-9140-A13, Broadcom, San Jose, CA) mounted under the wheel.

### Immunohistochemistry

Deeply anesthetized animals were transcardially perfused with 0.01 M PBS (pH 7.4) followed by 4 % paraformaldehyde. Brains were post-fixed for 6 h, cryoprotected with 30 % sucrose solution in M PBS, and serially cut in coronal sections (thickness: 40 - 50 µm) using a HM 450 Sliding Microtome (Thermo Fisher). Sections were counterstained with Hoechst (1:300, Sigma Aldrich, Milan, IT), mounted, and coverslipped with a DABCO [1,4-diazobicyclo-(2,2,2)octane]-based antifade mounting medium. Fluorescence images were acquired with a Leica SP5 inverted confocal microscope (Leica Microsystems, Milan, IT).

For the evaluation of the anatomical connections between VPM and S1bf, mice were perfused after 10 days from the injection and GRIN lens implantation. 50 µm thick coronal brain slices were cut, counterstained with Hoechst (1:300, Sigma Aldrich, Milan, IT), and mounted with an Antifade Mounting Medium (Vectashield, Burlingame, CA). Confocal images were acquired with a Nikon Eclipse scope (Nikon, Milan, IT).

### Simulations

#### Geometrical considerations

In Fig. 4a-h, neurons were simulated as spheres with an average radius *r_mean* which was estimated from recorded data (*r_mean* = 7.95 ± 2.33 µm, mean ± sd). Some variability was introduced in the neurons size, sampling it from a normal distribution with mean *r_mean* and standard deviation *r_sigma* = 1.31 (within the measured sd). At the center of each neuron a nuclear region was added, such that the spherical shell surrounding the nucleus had a width of variable size (*r_shell* randomly sampled from a normal distribution with mean = 4 µm and sd = 1 µm). The nuclear region did not express GCaMP6s and the fluorescence signal could be collected only from the spherical shell surrounding the nucleus. Simulated neurons were randomly placed in a volume of size 500 x 500 x 80 µm^3^ with no overlap between neurons up to the point that the volume was filled with cells or that neural density reached the value 83,100 ± 7,900 cells/mm^3 80^. The resolution of the spatial volume was 0.5 µm/pixel in the x and y direction, 1 µm/pixel in the z direction.

#### Neural activity

Neural spiking activity was simulated as the sum of Poisson processes. Each neuron was assigned with a mean spiking rate (*rho* = 0.4, arbitrary selected), following a binary synchronicity matrix with value 1 for neurons with common inputs (*common inputs probability* = 0.8, arbitrary selected). The activity of each neuron was the sum of an independent Poisson process and as many common Poisson processes as the neurons with shared variability due to common inputs. The spiking rate of the summed Poisson processes was the mean spiking rate for that neuron. Calcium activity and fluorescence traces were then generated using the equations in ^81^. An autoregressive model of order 1 (with parameter γ = 0.7) was selected to convolve spike trains into calcium activity. A model with supralinearities and Hill saturation was used to convert calcium activity into fluorescence intensity (model parameters: baseline = 0; single event amplitude = 1500; Hill saturation coefficient = 1; dissociation constant = 1; noise variance = 0.05).

#### Generation of fluorescence time series

The size and the resolution of the simulated FOV were set to 500 x 500 µm^2^ and 2.5 µm/pixel, respectively. The resolution was adjusted according to the changes in the magnification factor (estimated from experimental data Supplementary Fig. 4), obtaining a non-uniform resolution in the FOV.

To generate the synthetic t-series, we used the measurements experimentally obtained from corrected and uncorrected microendoscopes (Fig. 3). For corrected microendoscopes, the synthetic imaging focal surface was a spherical shell with curvature radius estimated from the measurements of fluorescent films (curvature radius: 400 μm) and the excitation volume was an ellipsoid resembling the aberration-corrected and experimentally measured PSF. For uncorrected microendoscopes, the synthetic imaging surfaces were two spherical shells with curvature radius estimated from the data (curvature radius: 265 μm and 2000 μm, respectively) and the excitation volume was an ellipsoid resembling the aberration-uncorrected and experimentally measured PSF. Excitation volumes were scanned along the imaging focal surface (or surfaces for uncorrected microendoscopes), such that their axial direction was always orthogonal to the imaging focal surface(s). All the voxels falling within the excitation volumes contributed to the signal of the corresponding pixel in the FOV, resulting in one of the following possible three conditions:

- If the pixel was in the edge of the FOV (radial distance > 250 µm), its signal was randomly sampled from a normal distribution, with mean and standard deviation estimated from experimental data. Dark noise mean was best fitted by a Gaussian mixture model (component 1: proportion = 0.37; mean = 137.48; sd = 48.96; component 2: proportion = 0.63; mean = 126.83; sd = 5.02), while the standard deviation of the dark noise depended on the dark noise mean in a linear way (p_0_ = −175.39, p_1_ = 1.57). The simulated dark noise was generated with the mean randomly sampled from the Gaussian Mixture Modeling (GMM) distribution and the standard deviation linearly dependent from the mean.
- If the pixel was in the central part of the FOV (radial distance ≤ 250 µm) but no neurons were within the excitation volume, the pixel signal was randomly sampled from a normal distribution with mean and standard deviation estimated from experimental data. The mean intensity of pixels that were neither in the edges nor belonging to ROIs were fitted using a lognormal distribution (mean = 5.43, sd = 0.36) and the best linear fit between the squared root of the mean intensity and the intensity sd was computed (p_0_ = −162.55, p_1_ = 18.28). Simulated noise in the FOV was generated as Gaussian noise with mean randomly sampled from the lognormal distribution and sd linearly dependent from the squared root of the mean.
- If the pixel was in the central part of the FOV (radial distance ≤ 250 µm) and at least one neuron was in the excitation volume(s), each voxel in the excitation volume(s) was assigned either Gaussian noise (estimated as in the previous condition) in case no neurons were in that voxel, or the fluorescence intensity of the neuron sampled by that voxel. In case a neuron was contained in a voxel, Gaussian noise was also added to the neuron signal. The mean of the added Gaussian noise was zero, while the sd was proportional to the square root of the mean intensity of the voxel, with the coefficients estimated from a linear fit between the square root of the mean intensity and the intensity standard deviation of pixels assigned to ROIs in experimental data (p_0_ = −132.44, p_1_ = 16.94). The activity of all the voxels falling within the excitation volume(s) was then averaged to obtain the pixel’s fluorescence intensity. The intensity of each pixel signal was finally modulated as a function of the radial position within the FOV, accordingly to the optical characterization of corrected and uncorrected microendoscopes using the radial intensity obtained imaging the subresolved fluorescent layer (Fig. 3).

#### Segmentation of simulated time series

We implemented an automated procedure for the segmentation of synthetic t-series, based on the ground truth spatial distribution of neurons in the FOV. We first associated to each imaged neuron its spatial footprint, which consists of all the pixels collecting signal from that neuron. This segmentation would be the ideal segmentation. However, for experimental data, the ground truth is not available neither to users segmenting the FOV nor to automated segmentation algorithms. For this reason, we modified the ideal segmentation to obtain a more realistic situation. We reasoned that very small ROIs are not likely to be detected and therefore we removed all the ROIs with few pixels. We tried different thresholds for the minimum number of pixels composing a ROI (n = 5,10,15 pixels). We observed that this parameter did not have an effect on the results of the comparison between corrected and uncorrected microendoscopes and therefore we set it to 5. We then reasoned that overlapping ROIs could be distinguished only if their overlap was not perfect and therefore we merged ROIs with high overlap. The overlap between two ROIs was defined as the fraction between overlapping pixels and total number of pixels of the smallest ROI. We merged ROIs with overlap larger than 70, 80 and 90 %. This parameter had no effect on the results of the comparison between corrected and uncorrected microendoscopes and we set it to 80 %. Ultimately, we considered that ROIs whose fluorescence signal had low SNR could not be discriminated from noise. We reasoned that a single event was sufficient to segment a ROI and we defined a peak-SNR as the ratio between the maximum peak in the fluorescence trace and the baseline noise (defined as the sd of those parts of the fluorescence trace with signal lower than the 25 % of the fluorescence distribution). We considered in the final segmentation only ROIs whose signal peak-SNR was higher than a threshold of 5, 10, 15, 20, 25, and 30 (Fig. 4).

## Statistics and analysis

### Statistics

Values are expressed as mean ± sem, unless otherwise stated; the number of samples (n) and p values are reported in the Figure legends or in the text. No statistical methods were used to pre-determine sample size. All recordings with no technical issues were included in the analysis and blinding was not used in this study. Statistical analysis was performed with MATLAB software (Mathworks, Natick, MA) and GraphPad Prism software (GraphPad Software, San Diego, CA). A Kolmogorov-Smirnov test was run on each experimental sample to test for normality and to test the equality of the three distributions in Fig. 6g. The significance threshold was always set at 0.05. When comparing two paired populations of data, paired Students’s *t*-test or paired Wilcoxon signed-rank test (Fig. 4c) were used to calculate statistical significance in case of normal and non-normal distribution, respectively. Unpaired Student’s *t*-test (Fig. 4k) and Mann-Whitney test (Figs. 4e,l and 7d,g) were used for unpaired comparisons of normally and non-normally distributed data, respectively. One-way analysis of variance (ANOVA) with Bonferroni post-hoc correction was used to compare the dependence of multiple classes from a single factor (Figs. 6e,f and 7e). Two-way ANOVA with Bonferroni post-hoc correction was used to compare the dependence of multiple classes from two factors (Fig. 6g). Two-way ANOVA with the interaction factor and with Bonferroni post-hoc correction was used in Fig. 4d. To fit linear data, linear regression was used. The significance of linear regression coefficients was assessed using a Student’s *t*-test (Figs. 4g,m and 7h,i). Pearson correlation coefficients were used to test the dependence between variables (Fig. 7c,f,k). The significance of the Pearson correlation coefficients was assessed using a Student’s *t*-test. All tests were two-sided, unless otherwise stated. Information theoretical analyses, NMF modules identification, modules characterization and SVM classification were performed on MATLAB software (Mathworks, Natick, MA) using available toolboxes ^82^ or custom written codes.

### Analysis of confocal images

Three mice were unilaterally injected with AAV-eGFP and red retrobeads after tissue aspiration and then implanted with the endoscope. Three mice were injected with AAV-GFP and red retrobeads without tissue aspiration and they were not implanted (controls). Three confocal images were acquired from fixed slices for each hemisphere at different focal planes (minimal distance between planes: 20 µm). Images in the red and green acquisition channels were blurred with a Gaussian filter (sigma = 2 µm) and binarized with a triangle thresholding method. S1bf was manually identified using anatomical cues form Hoechst labeling in each sample. To quantify the amount of preserved TC and CT connections within a given area of S1bf, we computed the fraction of pixels showing suprathreshold pixel intensity out of the total of pixel of the chosen area. A single value for each sample, obtained by averaging between confocal images of different FOVs, was used to run the one-tailed Mann-Whitney test for the different acquisitions channels (Fig. 5).

### Analysis of field distortion and calibration of the pixel size

A regular fluorescent grid spanning the FOV was imaged in order to evaluate the distortion in the FOV. The number of pixels necessary to span 10 µm in the x and y direction was measured as a function of the distance from the FOV centre. A magnification factor which varied along the radial directions was evaluated by computing the ratio between the measured number of pixels in the distorted (microscope objective coupled with GRIN lens based microendoscope) and undistorted (microscope objective alone) conditions. The estimated magnification factor (from x and y directions) was fitted using a quadratic curve (corrected: p_0_ = 0.76, p_1_ = −6.24e-04, p_2_ = 1.95e-05, norm of residuals = 1.78; Uncorrected: p_0_ =0.73, p_1_ = −2.91e-04, p_2_ = 8.11e-06, norm of residuals = 0.24). The magnification factor was used to correctly calibrate experimental measurements in Figs-3,4,6,7 and Supplementary Fig. 3.

To measure the PSF as a function of the radial position within the FOV, z-stacks of subresolved fluorescent beads (diameter: 100 nm) were taken at different distances from the optical axis. Intensity profiles obtained from sections in the x, y, z directions of the PSFs were fitted with Gaussian curves and their FWHM was defined as x, y, and z resolution, respectively. Lateral resolution was calculated as the average of x and y resolution. Axial resolution coincided with the z resolution. When, due to aberrations in the lateral portion of the FOV, the intensity profile in the z direction was better fitted with a sum of two Gaussian curves instead of a single one, the axial resolution was defined as the axial distance between the two peaks of the best fitting curves. For each group of measurements at a specific distance from the optical axis, outliers were identified using the Rout method (GraphPad Software, San Diego, CA) and excluded from the dataset. Mean and standard deviation of resolutions were plotted against radial distance (Fig. 3). Data were fitted with a symmetric quartic function to respect the cylindrical geometry of the optical system and the maximal FOV radial extent was determined as the radial distance at which the axial resolution fitting curve crossed a 10 µm threshold. PSF measurements were conducted using at least three eFOV-microendoscope for each type.

### Analysis of fluorescence t-series

Experiments in VPM were analyzed with a customized graphical user interface (GUI) in MATLAB (version R2017a; Mathworks, Natick, MA). The GUI enabled the loading of data saved from the microscope acquisition software, included the motion correction algorithm (NoRMCorre) described in ^83^, facilitated the manual segmentation of the FOV, and allowed the deconvolution of neural activity from the recorded fluorescent dynamics. ROIs were drawn by visualizing single frames or temporal projections of the FOV and, within the selected regions, only those pixels necessary to maximize the peak-signal-to-noise ratio (SNR) of the mean intensity were selected. Peak-SNR was defined as:

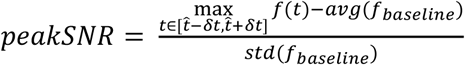

where *f*_*baseline*_ is the portion of intensity trace lower than the 25^th^ percentile of the intensity distribution. After segmentation, the GUI performed the deconvolution of normalized calcium activity from the fluorescence extracted from ROIs, by using the algorithm provided in ^84, 85^. The algorithm was based on the fit of the fluorescence activity with an autoregressive model. We used models of order 1 if the acquisition rate was low (< 2 Hz), otherwise order 2. At the end of the pre-processing, a structure containing all the extracted information (acquisition parameters, ROIs spatial footprints, ROIs fluorescence activity, deconvolved activity, and normalized calcium activity) was saved and used for the subsequent analyses.

For validation experiments in the hippocampus (Supplementary Fig. 6), temporal series recorded in the scanning configuration were imported into the open source ImageJ/Fiji software and movement correction was performed using the plugin Image Stabilizer. Calcium traces were analyzed using a custom code based on the open-source CellSort MATLAB toolbox ^86^. Briefly, the motion-corrected image stack was normalized and analyzed by principal components analysis (PCA) to find and discard dimensions that mainly represented noise. Principal components displaying a variance greater than noise were then analyzed with an iterative independent component analysis (ICA) algorithm to identify active cells. Manual validation of extracted traces was performed. Signals S_i_(t) were standardized as:

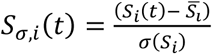

where 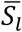 and σ(S_i_) are respectively the mean of the signal and its standard deviation.

### Analysis of synthetic and experimental time series

#### Quantification of ROIs number

For synthetic data in Fig. 4, the number of segmented ROIs was computed as a function of the probe type (uncorrected or corrected microendoscopes), of the peak-SNR value used in the automated segmentation procedure, and of the interaction between these two factor. For experimental data in Figs. 4,6,7, background intensity was not always uniform in the endoscopes FOV and in some regions no ROIs could be detected. In order to discount this factor, the count of the ROIs was normalized to the brighter part of the FOV, obtaining a measurement of the ROIs density (number of ROIs divided by the total bright area). To detect the dark background regions, the edges of the FOV (where mostly dark noise was collected) were used as intensity threshold. All the parts of the FOV with mean intensity lower than 85^th^ percentile of the threshold were discarded for the normalization of the ROIs count. The remaining part of the FOV was considered as bright area and used for the ROIs density analyses.

#### SNR of calcium activity

For both synthetic and experimental data, the SNR was defined as:

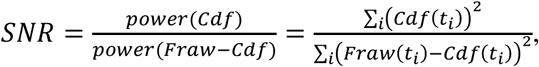

where *F*_*raw*_ and *Cdf* are the z-scored raw fluorescence intensity and deconvoluted calcium activity, respectively. SNR was evaluated to measure the quality of the extracted ROIs signal.

#### Correlation with ground truth activity

For synthetic data, we computed the correlation between the calcium activity of each segmented ROI and the ground truth calcium activity of neurons contributing to that ROI. In case more neurons were merged during the automated segmentation, we sorted the merged neurons for decreasing correlation with the corresponding ROI. We defined “source neuron” the neuron with highest correlation with the ROI (Fig. 4h left) and considered the correlation with the other merged neurons (only the second highest correlation is shown) as a measure of signals’ contamination between nearby neurons (Fig. 4h right).

#### Pairwise correlation

For each pair of nearby ROIs (distance between ROIs centers < 20 μm), we computed pairwise correlation of the extracted calcium activity as a function of the radial distance of the ROIs pair. We defined the radial distance of each pair as the distance between the FOV’s center and the center of the segment connecting the two ROIs’ centers. To measure changes in pairwise correlations exclusively caused by changes in imaging resolution, we normalized the pairwise correlations of nearby ROIs by subtracting the value of pairwise correlations between distant ROIs (distance > 60 μm) placed at the same radial distance. We then fitted the normalized pairwise correlation as a function of radial distance using linear regression.

### Analysis of behavioral parameters

Videos of whisker movements were binarized with the Ridge Detection plugin of ImageJ/Fiji in order to individuate pixels corresponding to whiskers. Videos were then processed in MATLAB (Mathworks, Natick, MA) to extract the whisker mean angle. To this aim, all whiskers were fitted with straight lines and for each frame the mean angle of all the lines was calculated with respect to the horizontal direction of the FOV. Once the mean angle of the imaged whiskers was calculated for each frame, this signal was processed with a moving standard deviation over a 400 ms window and a Gaussian filter over a 50 ms window. Whisking and no whisking periods were identified by binarizing the mean whisker angle with a temporal and amplitude threshold. While the temporal threshold was fixed at 200 ms, the amplitude threshold was extracted by manually identifying whisking periods in ∼1/10th of the full-length video and using this manual classification to find the best amplitude threshold with a ROC analysis. Temporal gaps between whisking periods shorter than 0.5 s were considered whisking periods, and linear interpolation was used to obtain the whiskers mean angle in frames in which less than 4 whiskers were detected.

For the analysis of pupil diameter, movies were analyzed with MATLAB (Mathworks, Natick, MA). Each frame was thresholded with the Otsu’s method and the region corresponding to the pupil was approximated with an ellipse. The length of the major axis of the ellipse was considered the pupil diameter. Linear interpolation was used for frames in which the pupil was not properly detected.

Detection of locomotion periods was performed using a threshold criterion on the wheel speed ^87^. The wheel speed signal was downsampled at 40 Hz and an instant was considered to be part of a locomotion epoch if it met the following conditions: *i)* instantaneous speed > 1 cm/s; *ii)* low-pass filtered speed (short pass filter at 0.25 Hz) > 1 cm/s; *iii)* average speed over 2 s windows > 0.1 cm/s. Temporal gaps between locomotion periods shorter than 0.5 s were considered periods of locomotion.

Four behavioural states were defined in Fig. 6: *i)* quiet (Q) when neither locomotion nor whisking were observed, *ii)* whisking (W), when whisking but no locomotion were observed; *iii)* locomotion (L), when locomotion but not whisking were detected; *iv)* whisking and locomotion (WL), when both locomotion and whisking were detected. L epochs were extremely rare (1.45 ± 0.75 % of total acquisition time, mean ± sem) and were not considered in the analysis. For the SVM analysis and for the NMF analysis (Figs. 6,7), we considered just two states: quiet (Q) and active (A), with A being the union of W and WL.

### Analysis of calcium signals across states

To compare the amplitude of calcium activity across behavioral states, the deconvolved activity of each ROI was averaged in each of the three states (Q, W and WL).

To measure whether calcium activity was further modulated by arousal, we discretized the pupil size in ten bins and measured the distribution of the average calcium activity for each behavioral state, separately.

### Information theoretic analysis

For information theoretical analyses, we used the MATLAB toolbox provided in ^82^. To compute whisking information encoded in the calcium signal of single ROIs, we computed the amount of mutual information ^88^ that the calcium signals carried about whether the animal was in a quiet (Q) or active (A) state. For each state, *n*_*T*_ time points, where *n*_*T*_ is the number of time points spent in the less frequent state, were randomly sampled without replacement. For each ROI then, the calcium activity in the selected time points was used to compute the information carried by that ROI regarding the state of the animal (the details were as follows: direct method, quadratic extrapolation for bias correction, number of response bins = 2). The amount of information was considered significantly different from 0 only if the real information value was larger than the 95^th^ percentile of the distribution of n = 500 shuffled information values (obtained after randomly shuffling the whisking state label). This procedure was repeated n = 100 times, randomly sampling different time points and the reported information values were computed as the average information encoded across different iterations.

We sorted ROIs according to their information content and fitted the distribution of information content of individual ROIs using a double exponential function, where the information carried by the *i*-th ROI was given by:

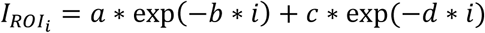

We used the R^2^ coefficient to assess the goodness of the fit. We computed the Pearson correlation coefficient to check for dependence between the information carried by individual ROIs and the ROIs radial distance (or the ROIs SNR). To test whether ROIs at different radial distances (or with different SNR) carried a different amount of information, we split the ROIs in two groups. ROIs with low radial distance (or SNR) where those ROIs whose radial distance (or SNR) was lower than the median of the radial distances (or SNR) distribution. ROIs with high radial distance (or SNR) where those ROIs whose radial distance (or SNR) was higher (or equal) than the median of the radial distances (or SNR) distribution.

To compute information carried by pairs of neurons, we considered only pairs of nearby neurons (distance between neurons < 20 μm). We computed the amount of synergistic information carried by the neuronal pair as a function of the pairwise correlation between neurons and as a function of the pair’s radial distance. We fitted the synergistic information using a linear function for synergistic and redundant pairs, separately.

To compute information about the whisking state from a large population of neurons, we first decoded the whisking state from the single trial population activity, and then we expressed the decoding performance as mutual information contained in the confusion matrix as in equation (11) of ^88^. We performed single trial population decoding as follows. A Gaussian kernel SVM was trained to classify the animal state (Q or A) by observing population activity ^89^. For each state, *n*_*T*_ time points were randomly sampled (without replacement) and split into two equal and balanced sets. One set was used as training set for the SVM, and on this data, a ten-fold cross validation was performed over a fixed grid for the SVM hyperparameters. The performance of the SVM were then tested using the test set. This procedure was repeated n = 100 times by randomly sampling different time points. The reported classification accuracy was the average information encoded across different iterations. To check whether an increase in the number of imaged ROIs led to a better classification of whisking state (Fig. 7e), we computed information for neuronal population of gradually increasing size. At first, we considered only the ROIs in the central portion of the FOV (distance from FOV center < ¼ of FOV radius) and then the other ROIs (distance steps = ¼ of FOV radius) were incrementally added for the training and testing phase of the SVM.

### Non-negative matrix factorization (NMF)

To compute NMF (Fig. 7), two states (Q and A) were considered. For each state, *n*_*T*_ time points were randomly sampled (without replacement) and split into two equal and balanced sets (a training set and a test set). The number of NMF modules was selected based on the ability of a linear discriminant analysis (LDA) classifier trained on the reduced data to predict the presence or absence of whisking ^90^. The dimensionality of the training set and the test set were at first reduced to k, with k ranging from 1 to the number of ROIs of the dataset. Then, for each factorization, the LDA classifier was trained on the training set to predict the behavioural state variable, and its performance was tested on the test set. The final dimension for the NMF was selected as the number of modules at which the first elbow in the performance plot (performance increase < 0.4%) was found. Then, the dimensionality of the entire dataset was reduced by computing the NMF with the selected number of modules. For each module in the obtained factorization, the following quantities were computed:

- Sparseness: 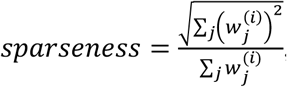, where 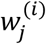 denotes the contribution of the j^th^ ROI to the *i*^th^ module. Sparseness values close to 1 indicate that few ROIs contribute heavily to the module, while sparseness values close to 0 indicate that the contribution of ROIs to the modules is more homogeneous.
- Whisking modulation index (WMI): 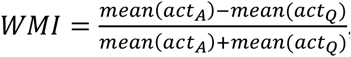, where *act*_*A*_ and *act*_*Q*_ denote the activation coefficients in each behavioral state. WMI > 0 indicates that the module’s activity is increased during A state, while WMI < 0 indicates that the module’s activity is reduced during A state.
- Spatial spread: we defined as spatial spread the shortest path that connected the ten ROIs with highest weights (or all the ROIs in case a module was composed by fewer ROIs).

To compare pairs of modules, we computed the following similarity measures:

- The Jaccard index is the fraction between the number of ROIs belonging to both modules and the total number of ROIs composing the two modules (without repetitions). It ranges between 0 and 1 and assumes value 0 if two modules do not share common ROIs, value 1 if two modules are composed by exactly the same ROIs. The Jaccard index does not take into account the weights of ROIs in the modules.
- Cosine similarity 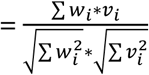, where _*i*_ and *v*_*i*_ represent the weight of the *i*-th ROI in each of the two modules. As for the Jaccard index, cosine similarity ranges between 0 and 1 and it takes value 0 if two modules do not share common ROIs. Contrarily to the Jaccard index, cosine similarity takes into account also the weights of the ROIs in the modules and it is 1 only if two ROIs are composed by exactly the same ROIs with equal weights.

## Acknowledgements

We thank M. Dal Maschio for discussion at an initial stage of the project, F. Nespoli for preliminary analysis, B. Sabatini and A. Begue for critical reading, V. Jayaraman, J. Akerboom, R. A. Kerr, D. S. Kim, L. L. Looger, K. Svoboda for GCaMPs. This work was supported by an IIT interdisciplinary grant and in part by ERC (NEURO-PATTERNS), NIH Brain Initiative (U01 NS090576, U19 NS107464, R01NS109961), FP7 (DESIRE), MIUR FIRB (RBAP11×42L), and Flag-Era JTC Human Brain Project (SLOW-DYN). CL acknowledges support from KAUST under baseline funding BAS/1/1064-01-01.

## Author contributions

A.A., A.S., S.B., C.M., and F.S. performed experiments. M.M., A.A., A.S., S.B., C.M., A.F. and performed analysis. A.A., A.S., and C.M. performed simulations and microendoscope fabrication. A.A. and A.B. performed microlens fabrication. V.P.R. developed polymeric materials and molding replication process. T.F. conceived and coordinated the project. T.F., C.L., and S.P. supervised the project. C.L. provided guidance in optical simulations and *eFOV*-microendoscope fabrication. S.P. provided advice on data analysis and simulations. All authors contributed to writing and approved the final version of the manuscript.

