## Supplementary Figures and Tables for "Extended field-of-view ultrathin microendoscopes for high-resolution two-photon imaging with minimal invasiveness in awake mice"

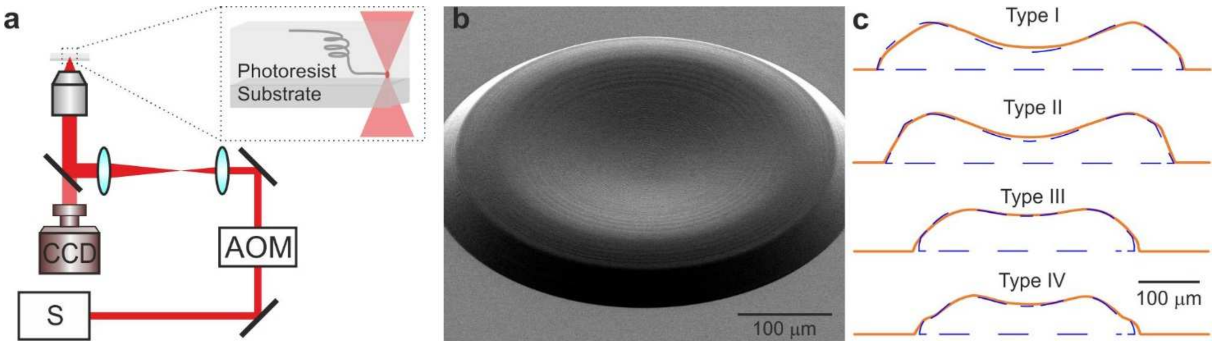


**Supplementary Fig. 1. Polymer lens fabrication using 3D micro-printing based on two-photon lithography. a,** Schematic of the optical set-up for two-photon lithography. AOM, AcoustoOptical Modulator; S, laser source; CCD, camera. **b,** Scanning electron microscope image of a corrective polymer lens. **c,** Blue dashed lines indicate the designed lens profile. Red lines represent the lens profile measured with a contact stylus surface profiler.

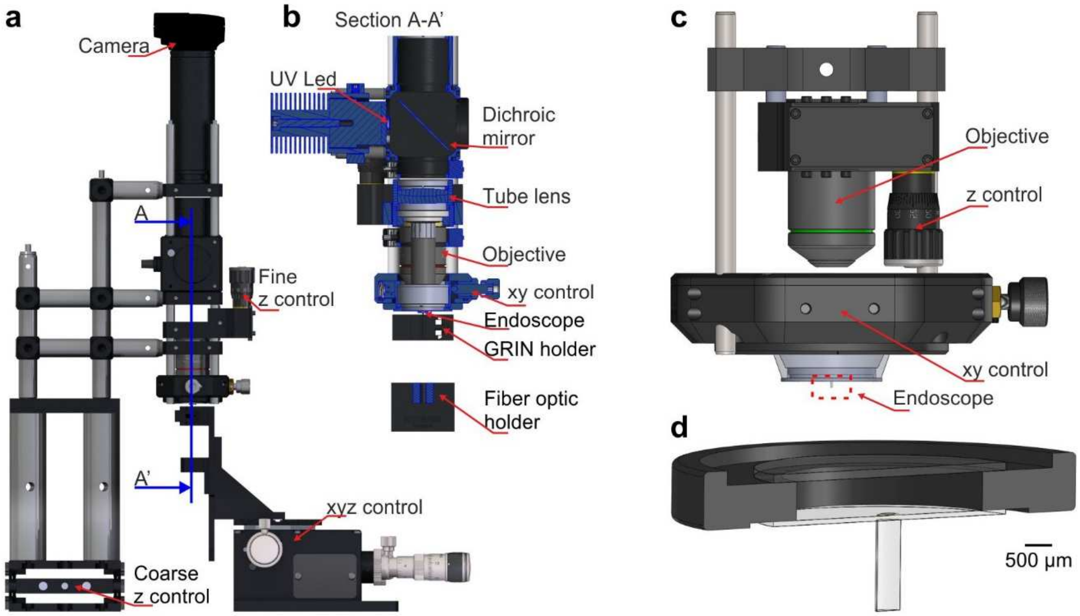


**Supplementary Fig. 2. Set-ups for the assembly and characterization of *eFOV*-microendoscopes. a,** Optomechanical stage used for microendoscope assembly. Red arrows indicate key components. The blue line indicates the plane used for the cross-section view shown at an expanded scale in **b**. **b,** Section of a portion in **a-a'** of the set-up shown in **a**. **c,** Schematic of the optomechanical assembly used for microendoscopic imaging. **d,** The *eFOV*-microendoscope assembly for implantation highlighted with the red dotted line in **c** is shown at an expanded scale.

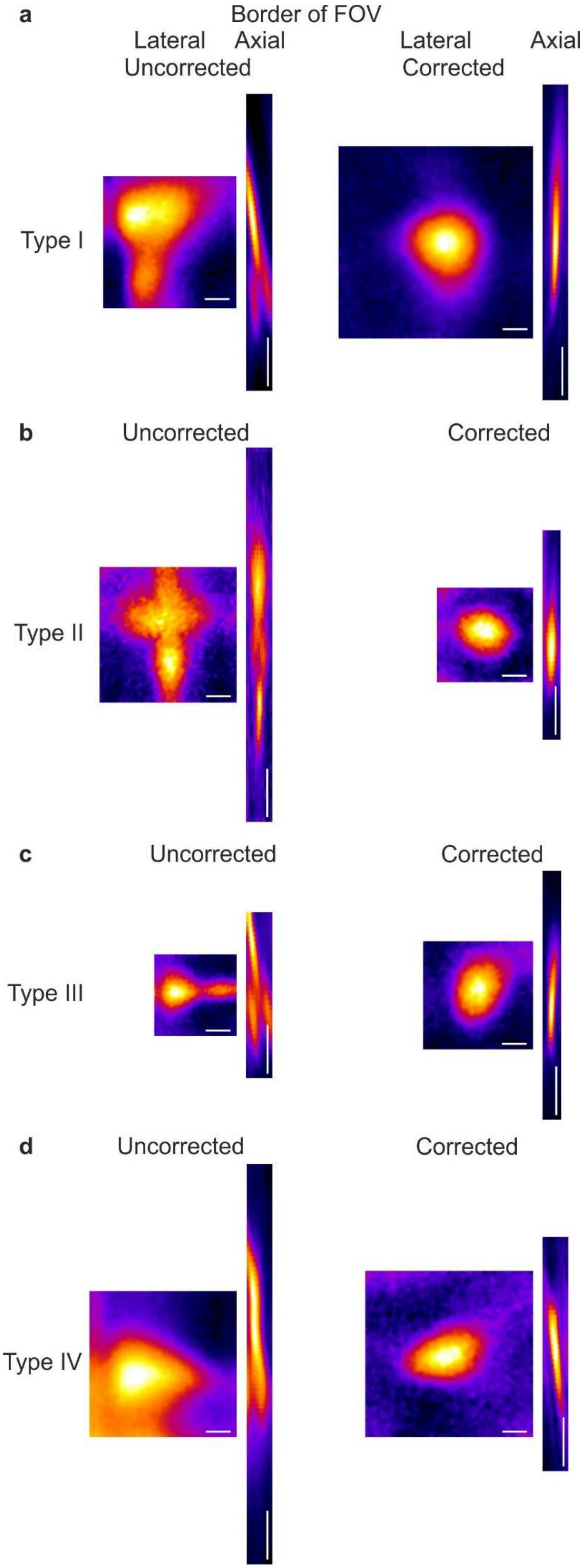
a
Border of FOV
Lateral
Axial
Uncorrected
Lateral
Axial
Corrected
Type I
b
Uncorrected
Corrected
Type II
c
Uncorrected
Corrected
Type III
d
Uncorrected
Corrected
Type IV

**Supplementary Fig. 3. Aberration correction improves the PSF in the peripheral portions of the FOV in *eFOV*-microendoscopes. a,** Lateral and axial projection of a z-stack of a subresolved fluorescent bead positioned at the border of the FOV and imaged with the uncorrected (left) and corrected (right) type I endoscopes. Scale bars: 1 µm (lateral), 10 µm (axial). **b,** Same as in **a** for type II endoscopes. **c,** Same as in **a** for type III endoscopes. **d,** Same as in **a** for type IV endoscopes. Please note that the slight tilt of the axial PSF with respect to the optical axis in corrected endoscopes reflects the curvature of the distal conjugated optical plane, as visible in Figs 1,3 (non telecentric optical system).

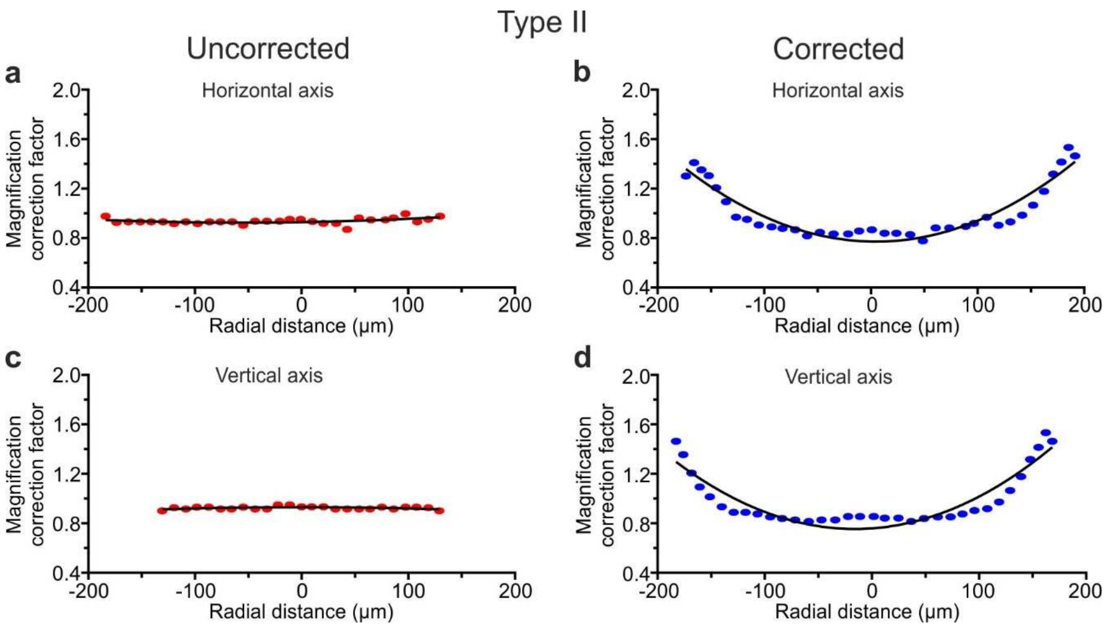


**Supplementary Fig. 4. Field curvature in *eFOV*-microendoscopes. a,b,** Magnification correction factor in the horizontal direction as a function of the radial position in uncorrected (**a**) and corrected (**b**) type II *eFOV*-microendoscopes. Plots show values obtained from one representative measurement. Dots represent experimental data. Black lines indicate quadratic polynomial fit. **c,d,** Same as in (**a,b**) for the vertical direction.

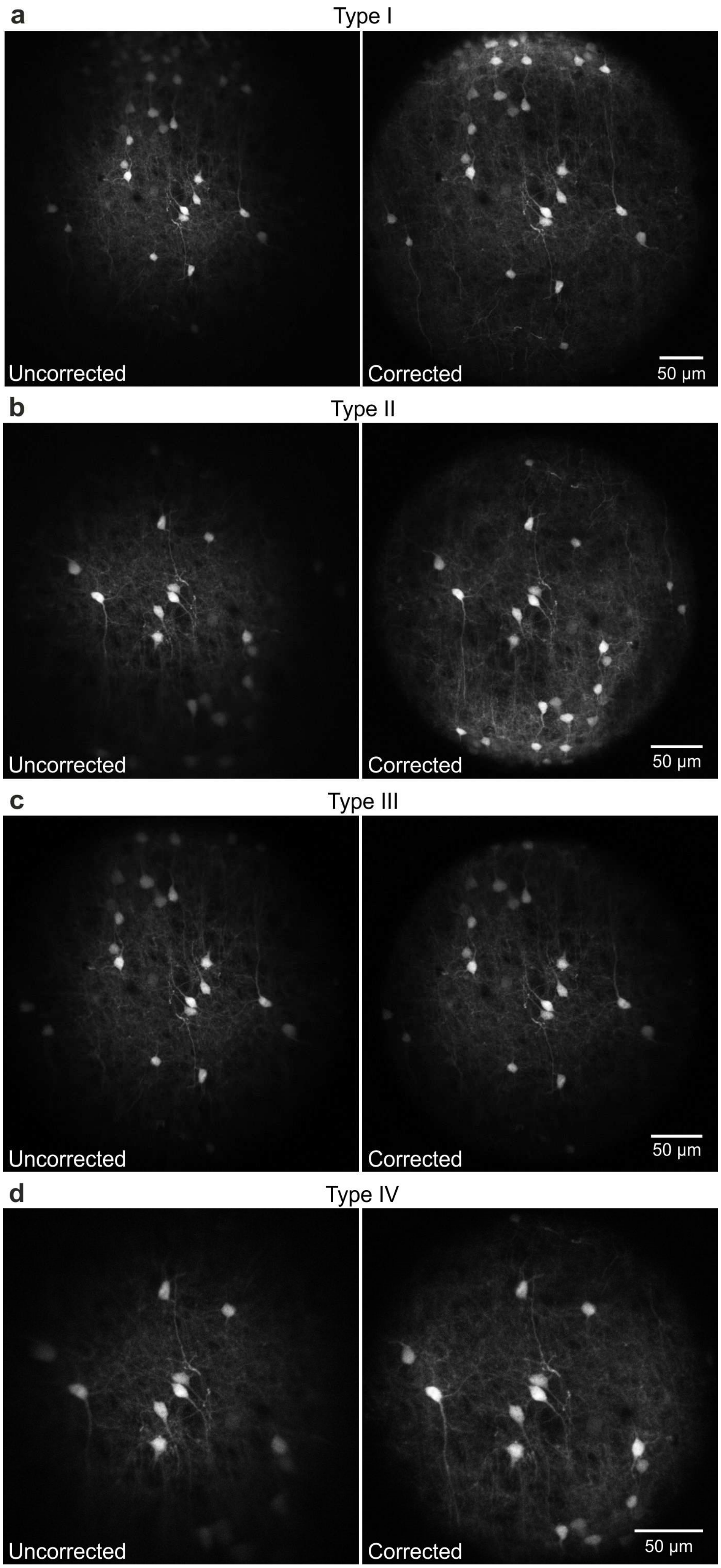
a
Type I
Uncorrected
Corrected
50 μm
b
Type II
Uncorrected
Corrected
50 μm
c
Type III
Uncorrected
Corrected
50 μm
d
Type IV
Uncorrected
Corrected
50 μm

**Supplementary Fig. 5. Corrected microendoscopes have extended effective FOV. a-d,** Representative images of fixed cortical tissue expressing eGFP in neuronal cells acquired with type I (**a**), type II (**b**), type III (**c**), and type IV (**d**) microendoscopes without (uncorrected, left panels) and with (corrected, right panels) corrective lenses.

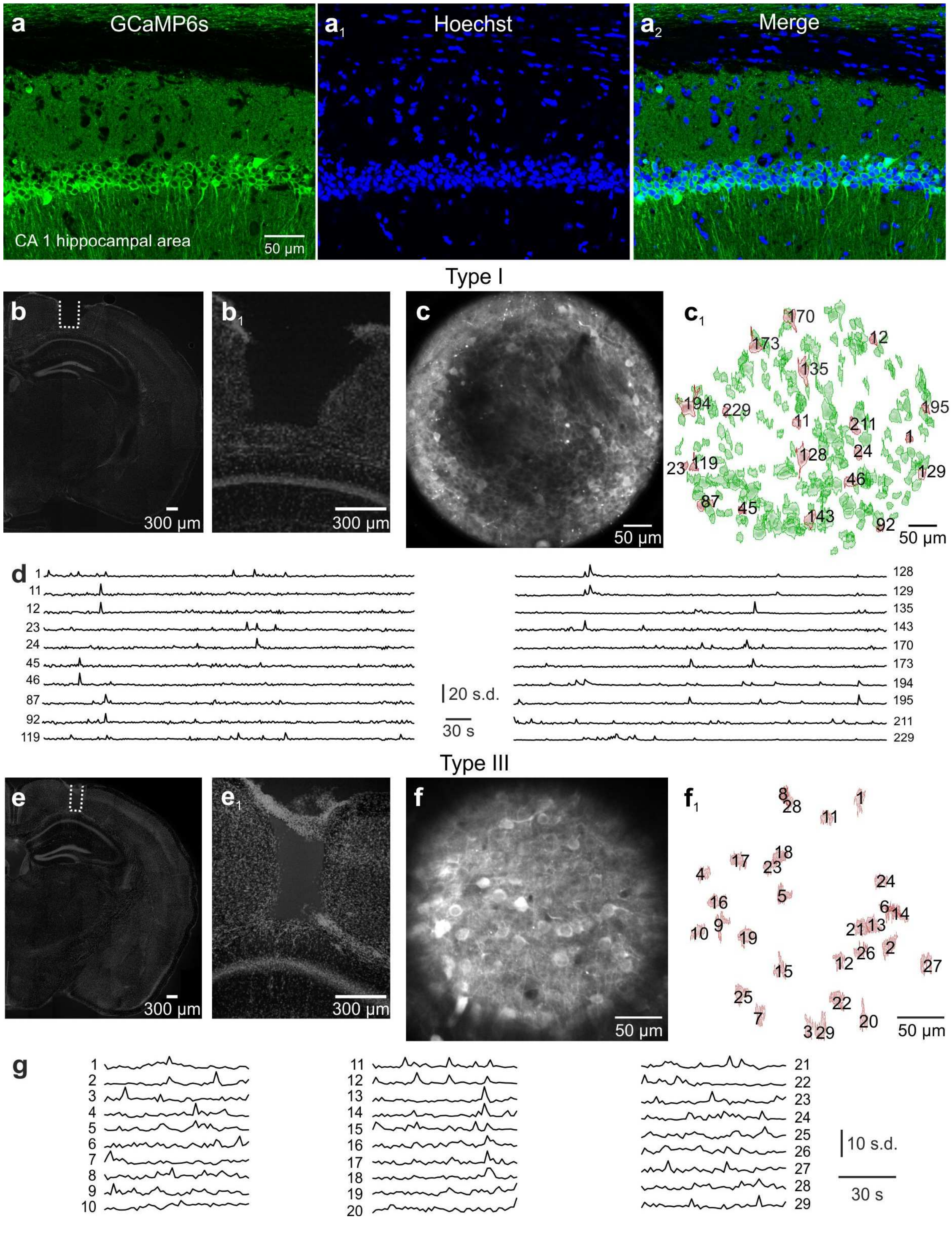


**Supplementary Fig. 6. Hippocampal imaging with type I and type III *eFOV*-microendoscopes. a-a$_2$,** Confocal images of hippocampal CA1 neurons expressing GCaMP6s (**a**). Nuclei were counterstained with Hoechst (**a**$_1$). Images are merged in (**a**$_2$). Scale bar in **a** applies to **a**$_1$-**a**$_2$. **b-b$_1$,** Confocal images showing coronal slices from a mouse implanted with type I *eFOV*-microendoscopes. Slices were counterstained with Hoechst. The probe track is highlighted with the white dotted line in **b** and shown at a higher magnification **b**$_1$. **c-c$_1$,** GCaMP6s expressing CA1 hippocampal neurons recorded by two-photon imaging using type I *eFOV*-microendoscopes in an anesthetized mouse. ROIs are identified in **c**$_1$. **d,** Fluorescence signals over time for the red ROIs shown in **c**$_1$. **e-e$_1$,** Same as in **b**-**b**$_1$ but for type III *eFOV*-microendoscopes. **f-f$_1$,** Same as in **c**-**c**$_1$ but for type III *eFOV*-microendoscope. **g,** Fluorescence signals over time for the ROIs displayed in **f**$_1$.

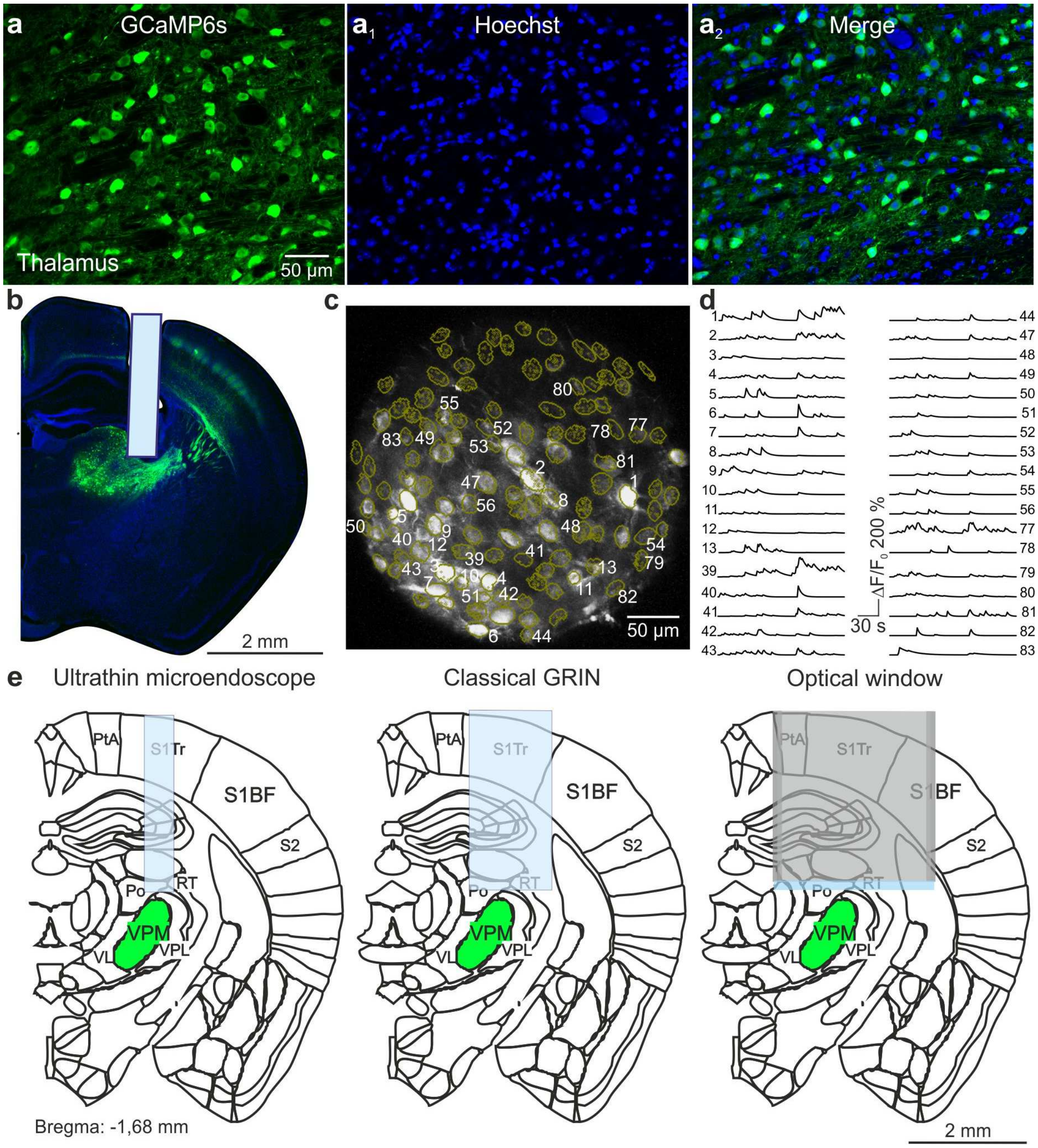


**Supplementary Fig. 7. VPM imaging with type II *eFOV*-microendoscopes. a-a$_2$,** Confocal images of thalamic neurons expressing GCaMP6s (**a**). Nuclei were counterstained with Hoechst (**a**$_1$). Images are merged in (**a**$_2$). Scale bar in **a** applies to **a**$_1$-**a**$_2$. **b,** Confocal image showing a coronal slice from a mouse implanted with a type II *eFOV*-microendoscope. The slice was counterstained with Hoechst. **c,** GCaMP6s-expressing VPM neurons recorded using type II *eFOV*-microendoscopes in an awake head-restrained mouse. ROIs are indicated in yellow. **d,** Fluorescence signals over time for some of the ROIs displayed in **c**. **e,** Schematic showing the implantation of an ultrathin microendoscope (left), of a larger cross-section GRIN lens (middle), and of an optical window (right) for VPM imaging. S1BF = primary somatosensory, barrel field; S1Tr = primary somatosensory, trunk region; PtA = parietal association region; S2 = second somatosensory cortex; VPM = ventral posteromedial thalamic nucleus; RT = reticular thalamic nucleus; Po = posteromedian thalamic nucleus; VPL = ventral posterolateral thalamic nucleus; VL = ventrolateral thalamic nucleus.

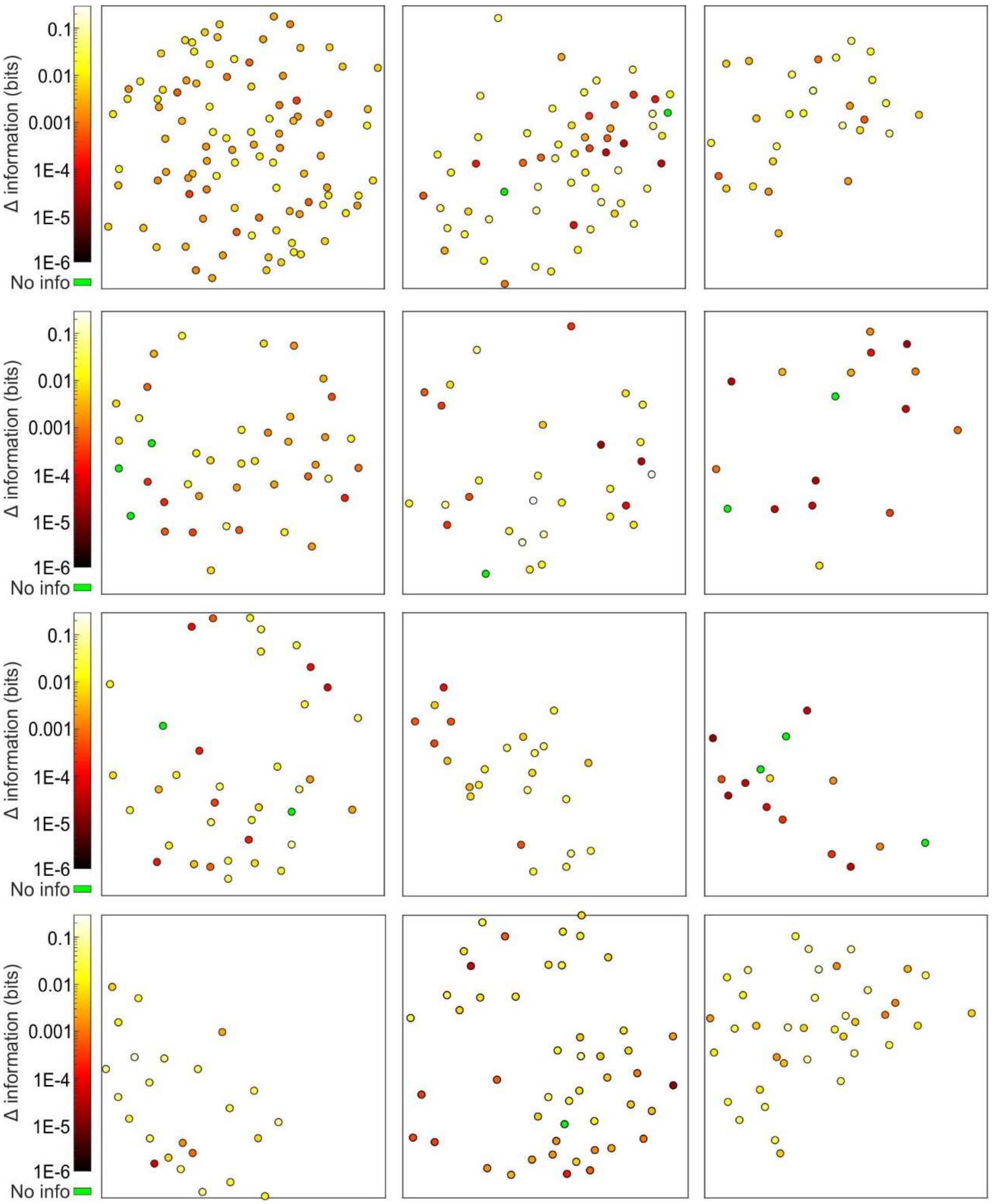


**Supplementary Fig. 8. Cell-specific encoding of behavioral state-dependent information in distributed VPM subnetworks.** Spatial map of neurons encoding whisking information in 12 out of the total of 24 analyzed time series. The pseudocolor scale shows significantly informative neurons (see Methods).

| | c | k | $\alpha_1$ | $\alpha_2$ | $\alpha_3$ | $\alpha_4$ | $\alpha_5$ | $\alpha_6$ | $\alpha_7$ | $\alpha_8$ |
|---|---|---|---|---|---|---|---|---|---|---|
| **Type I** | -2.6E-01 | -1.7E+00 | 8.6E-01 | -5.3E+01 | 6.0E+03 | -2.77E+05 | 7.3E+06 | -8.9E+07 | 2.5E+08 | 2.2E+09 |
| **Type II** | 1.0E+03 | -2.4E+03 | -1.5E+00 | 2.9E+01 | -4.6E+01 | -7.69E+03 | 2.0E+05 | 7.8E+06 | -1.7E+08 | -3.9E+08 |
| **Type III** | -5.0E-01 | 8.2E+00 | 1.5E+00 | -1.4E+02 | 3.7E+04 | -3.92E+06 | 2.2E+08 | -5.6E+09 | 3.3E+10 | 6.3E+11 |
| **Type IV** | -2.4E-01 | -1.2E+00 | 9.2E-01 | -1.3E+02 | 4.4E+04 | -4.28E+06 | 2.2E+08 | -5.6E+09 | 3.5E+10 | 4.9E+11 |

**Supplementary Table 1. Parameters for the fabrication of corrective lenses.** Coefficients used in equation (*1*) (see Methods) for the aspherical corrective lenses used in type I-IV *eFOV*-microendoscopes.

| Type | On axis $FWHM_{x,y}$ (μm) $n \geq 8$ | On axis $FWHM_z$ (μm) $n \geq 8$ | Effective FOV radius (μm) | Fold increase in FOV area |
|---|---|---|---|---|
| I | uncor.: 0.91 ± 0.01<br>cor.: 0.94 ± 0.02<br>p = 0.1607 | uncor.: 9.54 ± 0.18<br>cor.: 8.85 ± 0.12<br>p = 0.0026 | uncor.: 74<br>cor.: 133 | 3.2 |
| II | uncor.: 0.80 ± 0.01<br>cor.: 0.87 ± 0,02<br>p = 0.0064 | uncor.: 8.91 ± 0.15<br>cor.: 8.87 ± 0,11<br>p = 0.7916 | uncor.: 51<br>cor.: 156 | 9.4 |
| III | uncor.: 0.88 ± 0.02<br>cor.: 0.89 ± 0.00<br>p = 0.4298 | uncor.: 7.72 ± 0.07<br>cor.: 7.45 ± 0.16<br>p = 0.1861 | uncor.: 62<br>cor.: 154 | 6.2 |
| IV | uncor.: 0.78 ± 0.02<br>cor.: 0.89 ± 0.03<br>p = 0.0041 | uncor.: 8.86 ± 0.24<br>cor.: 8.32 ± 0.29<br>p = 0.1583 | uncor.: 34<br>cor.: 69 | 4.1 |

**Supplementary Table 2. Spatial resolution and effective FOV of *eFOV*-microendoscopic probes.** Value are reported as average ± sem. For statistical comparison of uncorrected (uncor.) *vs* corrected (cor.) microendoscopes, Student's *t*-test was used.

| Pupil diameter range | 0-0.1 | 0.1-0.2 | 0.2-0.3 | 0.3-0.4 | 0.4-0.5 | 0.5-0.6 | 0.6-0.7 | 0.7-0.8 | 0.8-0.9 | 0.9-1.0 |
|---|---|---|---|---|---|---|---|---|---|---|
| Q vs W | p = 1 | p = 1 | p = 1 | p = 1 | p = 1 | p = 1 | p = 0.7 | p = 1 | p = 1 | p = 1 |

| | | | | | | | | | | |
|---|---|---|---|---|---|---|---|---|---|---|
| Q vs WL | p = 1 | p = 0.04 | p = 7E-5 | p = 2E-6 | p = 2E-4 | p = 0.7 | p = 1 | p = 0.009 | p = 2E-6 | p = 2E-6 |
| W vs WL | p = 1 | p = 0.8 | p = 0.9 | p = 0.002 | p = 0.04 | p = 0.2 | p = 1 | p = 0.8 | p = 2E-6 | p = 2E-6 |

**Supplementary Table 3. Statistical comparisons of behavior state distributions as a function of pupil diameter.** For the statistical comparison of Q, W, and WL state distributions in each range of pupil diameter, a two-way ANOVA with Tukey-Kramer *post hoc* correction was performed.
